# Complementary task representations in hippocampus and prefrontal cortex for generalising the structure of problems

**DOI:** 10.1101/2021.03.05.433967

**Authors:** Veronika Samborska, James Butler, Mark Walton, Timothy E.J. Behrens, Thomas Akam

## Abstract

Few situations in life are completely novel. We effortlessly generalise prior knowledge to solve novel problems, abstracting common structure and mapping it onto new sensorimotor specifics. Here we trained mice on a series of reversal learning problems that shared the same structure but had different physical implementations. Performance improved across problems, demonstrating transfer of knowledge. Neurons in medial prefrontal cortex (mPFC) maintained similar representations across multiple problems, despite their different sensorimotor correlates, whereas hippocampal (dCA1) representations were more strongly influenced by the specifics of each problem. Critically, this was true both for representations of the events that comprised each trial, and those that integrated choices and outcomes over multiple trials to guide subjects’ decisions. These data suggest that PFC and hippocampus play complementary roles in generalisation of knowledge, with the former abstracting the common structure among related problems, and the latter mapping this structure onto the specifics of the current situation.

## 1 Introduction

When we walk into a new restaurant, we know what to do. We might find a table and wait to be served. We know that the starter will come before the main, and when the bill arrives, we know it is the food we are paying for. This is possible because we already know a lot about how restaurants work, and only have to map this knowledge onto the specifics of the new situation. This requires that the common structure is abstracted away from the sensorimotor specifics of experience, so it can be applied seamlessly to new but related situations.

Such abstraction has been variously described as a schema (in the context of human behaviour^1^ and memory research^2,3^), learning set^4^ (in the context of animal reward-guided behaviour), transfer learning^5^ and meta-learning^6^ (in the context of machine learning). We have little understanding of how the necessary abstraction is achieved in the brain, or how abstract representations are tied to the sensorimotor specifics of each new situation. However, recent data suggest that interactions between frontal cortex and the hippocampal formation play an important role^7^. Neurons^8,9^ and fMRI voxels^10,11^ in these brain regions form representations that generalise over different sensorimotor examples of tasks with the same structure, and track different task rules embedded in otherwise similar sensory experience^12,13^.

The involvement of these regions in abstraction is also of interest from a theoretical perspective. Both frontal cortex^14–17^ and hippocampus^18–27^ have been hypothesized to represent task states and the relationships between them. It has not been clear what distinguishes the representations in these regions, but some insight might be gained by considering hippocampal representations underlying spatial cognition. In rodent hippocampus, place cells are specific to each particular environment^28–30^, but firing patterns in neighbouring entorhinal cortex (including grid cells) generalise across different environments – that is, they are abstracted from sensorimotor particularities^31–35^. Similarly, there is evidence that mPFC representations of spatial tasks generalise across different paths^36–38^.

One possibility is that, as in space, abstracted or schematic representations of tasks in cortex might be flexibly linked with the sensorimotor characteristics of a particular environment to rapidly construct concrete task representations in hippocampus, affording immediate inferences^39,40^. Indeed, hippocampal manipulations appear particularly disruptive when new task rules must be inferred, either at the beginning of training^41^ or when task contingencies change^42,43^.

To probe cortical and hippocampal contributions to generalisation, we developed a novel behavioural paradigm where we presented mice with a series of problems with the same abstract structure (probabilistic reversal learning), but different physical instantiations, and hence different sensorimotor correlates. We recorded single units in medial prefrontal cortex (mPFC) and hippocampus (dCA1) across multiple physical port layouts in each recording session. We examined neuronal representations both of the individual elements of each trial, and of the cross-trial learning that controlled animals’ choices. Both mPFC and dCA1 representations of trial events were low dimensional, i.e. a small set of temporal patterns of activity, corresponding to tuning for particular trial events, explained a large fraction of variance in both regions. However, they differed with respect to how these representations generalised across problems. In mPFC, the same neurons tended to represent the same events across problems, irrespective of the sensorimotor particulars of the current problem. By contrast, although the same events were represented by hippocampus in each problem, the specific neurons that represented a given event differed in each problem. Both hippocampus and prefrontal cortex also contained representations of animals’ current policy that integrated events over multiple trials. These policy representations were again abstract in prefrontal cortex but tied to sensorimotor specifics in hippocampus.

## 2 Results

### Mice generalise knowledge between structurally equivalent problems

Subjects serially performed a set of reversal learning problems which shared the same structure but had different physical layouts. In each problem, every trial started with an ‘initiation’ nosepoke port lighting up. Poking this port illuminated two ‘choice’ ports, which the subject chose between for a probabilistic reward (Figure 1A). Once the subject consistently (75% of trials) chose the high reward probability port, reward contingencies reversed (Figure 1B). Once subjects completed ten reversals on a given port layout (termed a ‘problem’), they were moved onto a new problem where the initiation and choice ports were in different physical locations (Figure 1C). All problems therefore shared the same trial structure (initiate in the illuminated poke, then choose between the two illuminated pokes) and a common abstract rule (one port has high and one low reward probability, with occasional reversals), but required different motor actions due to the different port locations. In this phase of the experiment, problem switches occurred between sessions, and subjects completed ten different problems.

**Figure 1:**
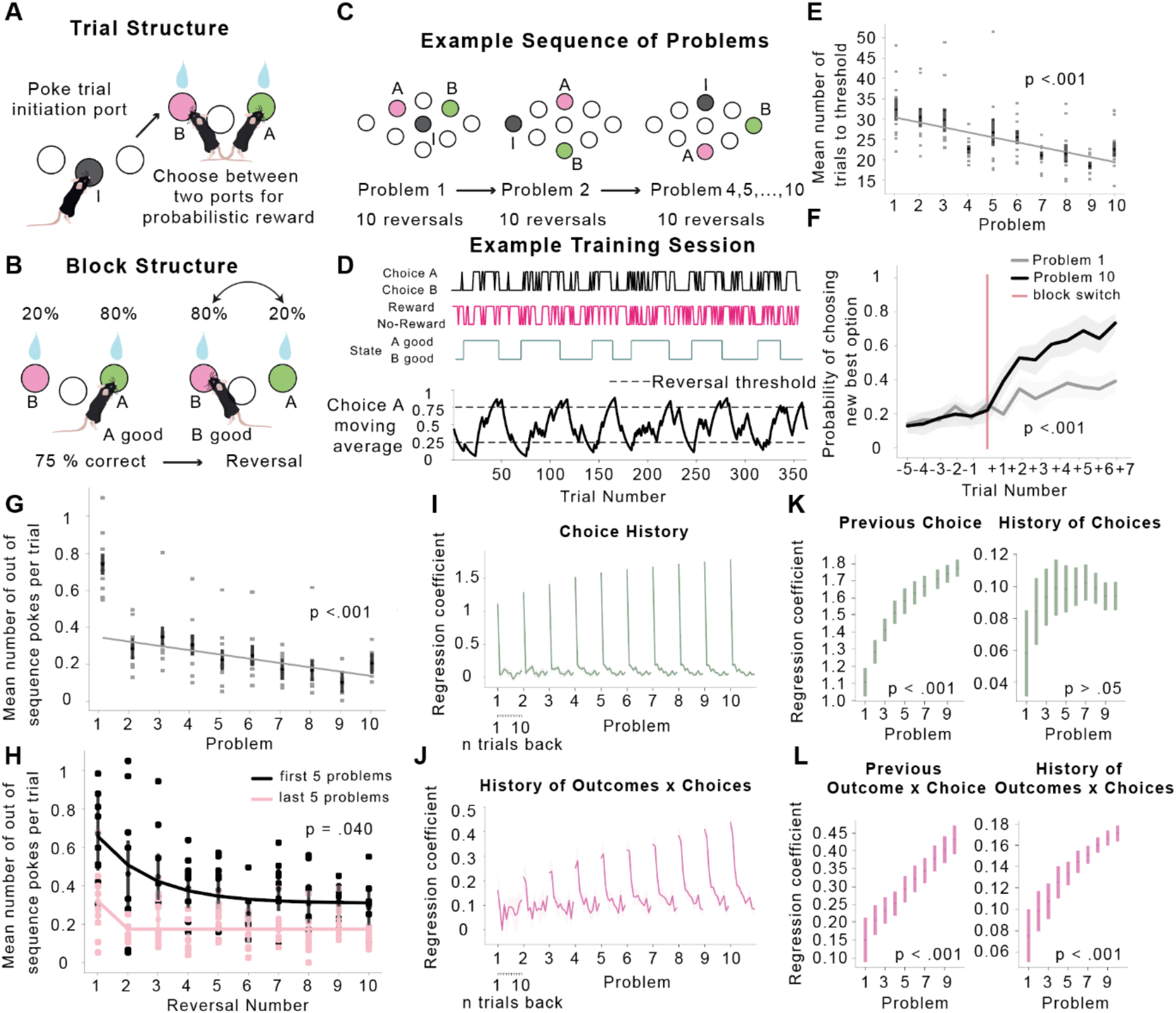
Transfer learning in mice. **A)** Trial structure of the probabilistic reversal-learning problem. Mice poked in an initiation port (grey), then chose between two choice ports (green and pink) for a probabilistic reward. **B)** Block structure of the probabilistic reversal-learning problem. Reward contingencies reversed after the animal consistently chose the high reward probability port. **C)** Example sequence of problems used for training, showing different locations of the initiation (I) and two choice ports (A & B) in each problem. **D)** Example behavioural session late in training in which the animal completed 12 reversals. Top subpanels show animals’ choices, outcomes they received, and which side had high reward probability; bottom panel shows exponential moving average of subjects’ choices (tau=8 trials). **E)** Mean number of trials following a reversal taken to reach the threshold to trigger the next reversal, as a function of problem number. **F)** Probability of choosing the new best option (the choice that becomes good after the reversal) on the last 5 trials before the reversal and the first 7 trials after the reversal split by the first problem and the last problem. The p-value refers to the difference between the slopes after the reversal point in early and late training. **G)** Mean number of pokes per trial to a choice port that was no longer available because the subject had already chosen the other port, as a function of problem number. **H)** Mean number of pokes per trial to a choice port that was no longer available as a function of reversal number on the first 5 problems and the last 5 problems during training. The p-value refers to the difference in the log of the time constants from fitted exponential curves in early and late training. **I, J)** Coefficients from a logistic regression predicting current choices using the history of previous choices **(I)**, outcomes (not shown) and choiceoutcome interactions **(J)**. For each problem and predictor, the coefficients at lag 1-11 trials are plotted. **K, L)** Coefficients for the previous trial (lag 1, left) and average coefficients across lags 2-11 (right), as a function of problem number. Error bars on all plots show mean ± SEM across mice.

We first asked whether subjects showed evidence of generalising the abstract problem structure (one port is good at a time, with reversals) to new problems (Figure 1B). Mice took fewer trials to reach the 75% correct threshold for triggering a reversal within each problem (*F*_(9,72)_ = 3.23, *p* = .002; Supplementary Figure 1A), and crucially also across problems (*F*_(9,72)_ = 3.88, *p* <.001; Figure 1E), consistent with generalising knowledge of this abstract structure. Improvement across problems in a subject’s ability to track the good port might reflect an increased ability to integrate the recent history of outcomes and choices across trials. To assess this, we fit a logistic regression model predicting subjects’ choices using the choices, outcomes, and choice-outcome interactions over the past history of trials. Across problems, the influence of both the most recent (*F*_(9,72)_ = 5.50, *p* <.001; Figure 1I,J) and earlier (*F*_(9,72)_ = 4.33, *p* =.001; Figure 1I,J) choice-outcome interactions increased. Subjects’ choices were also increasingly strongly influenced by their previous choices (*F*_(9,71)_ = 11.18, *p* <.001; Figure 1G,H), suggesting a decrease in spontaneous exploration with learning.

We also looked at whether subjects showed evidence of generalising the trial structure (initiate then choose; Figure 1A) across problems, by assessing how often they made nose pokes that were inconsistent with this sequence (i.e., pokes to the alternative choice port after having made a choice, instead of going straight back to initiation). Mice made fewer such out-of-sequences pokes across reversals within each problem (*F*_(9,72)_ = 5.14, *p* <.001; Supplementary Figure 1B), but importantly also across problems (*F*_(9,72)_ = 15.78, *p* <.001; Figure 1G). This improvement in knowledge of the trial structure was not just driven by animals’ poor performance on the first problem but continued to improve throughout training (*F*_(9,72)_ = 3.31, *p* =.003). To assess whether this improvement might be driven simply by animals learning to follow the light cues that signalled trial stages (and problem switches), we examined behaviour on ‘forced choice’ trials. On these trials only one of the choice ports was eliminated and the animals needed to select it before progressing with the task. Animals ignored the light and chose the best of the two choice ports, demonstrating that their behaviour was driven by their beliefs about where the reward was rather than port illumination (Supplementary Figure 2I, J).

A closer inspection of these data suggests that animals’ improvement across problems is consistent with meta-learning (or ‘learning to learn’). The definition of meta-learning is that when the animal needs to learn something it learns it faster each time. In line with this, early in training mice learnt the new poke sequences necessary to execute trials on new problems gradually, with the number of out-of-sequence pokes decreasing over many reversals, suggesting instrumental learning. However, at the end of the training they acquired the poke sequence in a single reversal suggesting they ‘learnt how to learn’ the new poke sequence (*t* _(17)_ = 2.45, *p* = .003, Figure 1H). Similarly, animals adapted to reversals faster at the end of training compared to beginning of training (*t* _(17)_ = 5.29, *p* < .001, Figure 1F). Therefore, they had also ‘learnt how to learn’ from reward.

These data suggest that mice learned to generalise both the block and trial structure across problems. We next searched for evidence of neural representations that abstracted the problem structure away from its physical details, allowing generalisation of knowledge.

### Abstract and problem-specific representations of trial events by PFC and CA1 neurons

We recorded single units from dorsal CA1 (345 neurons, n=3 mice, 91 to 162 neurons per mouse) and medial prefrontal cortex (mPFC, 556 neurons, n=4 mice, 117 to 175 neurons per mouse; Supplementary Figure 3, Figure 2) using electrophysiology. For recording sessions, we modified the behavioural task such that changes from one reversal-learning problem to the next occurred within a session, with the transition to the next problem triggered once subjects had completed four reversals on the current problem, up to a maximum of three problems in one session. Subjects adapted well to this change and in most recording sessions performed at least four reversals in three different layouts of the reversal-learning problem, allowing us to track the activity of individual units across problems (Figure 2B). Cross-problem learning reached asymptote prior to starting recordings, i.e., during recording sessions mice no longer showed improvement across problems (Supplementary Figure 2) and there were no differences in behavioural performance between CA1 and PFC animals (Supplementary Figure 2C, F).

**Figure 2:**
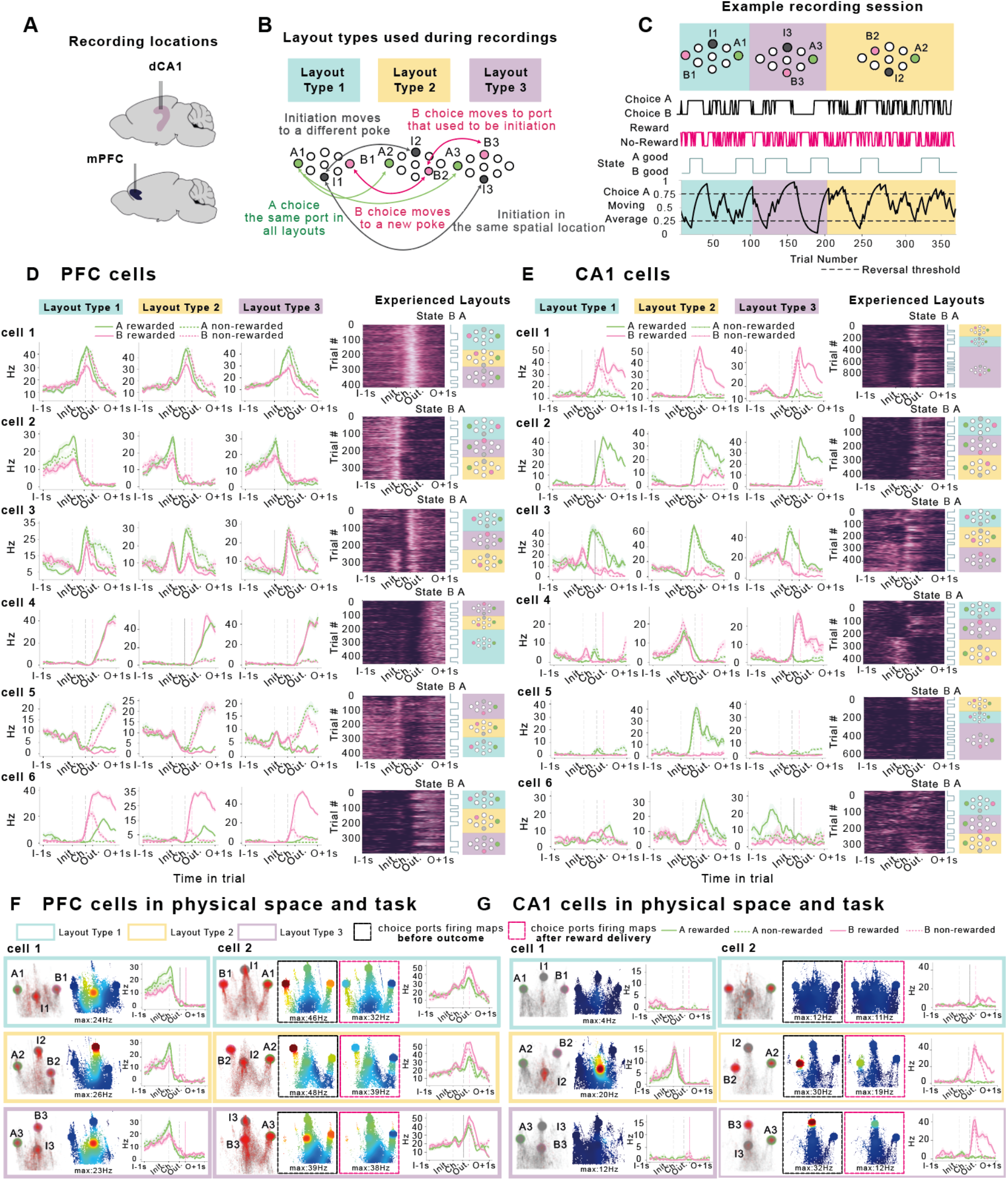
Recording units across multiple problems in a single session. **A)** Silicon probes targeting hippocampal dorsal CA1 and medial PFC were implanted in separate groups of mice. **B)** Diagram of problem layout types used during recording sessions. **C)** Example recording session in which a subject completed four reversals in each of three problems. Top panel shows the ports participating in each problem colour-coded by layout type. Bottom panel shows the exponential moving average of choices, with the choices, outcomes and reversal blocks shown above. **D) Example PFC neurons**. Cell 1 in PFC fired selectively to both choice ports (but not initiation) in each problem, even though the physical location of the choice ports was different both within and across problems. Cell 2 fired at the initiation port in every problem, even when its physical location changed. Cell 3 fired at B choice ports in all problems, but also gained a firing field when initiation port moved to the previous B choice port (showing PFC does have some port-specific activity). Cell 4 responded to reward at every choice port in every problem. Cell 5 responded to reward omission, and had high firing during the ITI. Cell 6 responded to reward at B choice port (that switched location) in each problem. **E) Example CA1 neurons**. Some CA1 cells also had problem general firing properties (cell 1 and 2). Cell 1 fired at B choice that switched physical location between problems. Cell 2 responded to the same port in all problems and modulated its firing rate depending on whether it was rewarded or not. Cell 3 fired at the same port in all layouts. Cell 4 switched its firing preference from initiation to B choice that shared physical locations, analogous to ‘place cells’ firing at a particular physical location. This port selectivity was more pronounced in CA1 than PFC (Supplementary Figure 10). Cell 5 and 6 ‘remapped’ - showing interactions between problem and physical port. Cell 5 fired at a given port in one layout but not when the same port was visited in a different layout. Cell 6 fired at choice time at a given port in one layout and changed its preferred firing time to pre-initiation in a different layout. In all plots average firing rates are arranged by layout type 1,2 and 3 but the order in which they were experienced is plotted on Experienced Layouts sub-panel. Shaded regions for firing rate indicate SEM across trials **F) Example PFC neurons in physical space and behavioural task.** For each cell, left panels show trajectories of animals’ nose (grey) and locations where spikes occurred (red) in a 2D space corresponding to the view of a camera positioned above the box looking at the ports, affine transformed to correct for the oblique view of the ports. Middle panels show firing rate heat maps in this same 2D space. Right panels show average firing rates across the trial for each trial type. Layout Types are indicated by the colour of boxes (blue, yellow, and purple). Cell 1 fired at the initiation port in every problem even when its physical location changed. Cell 2 fired at all choice ports in all problems. For choice port selective cells (PFC and CA1 cell 2), we split the firing rate maps by whether the within-choice-port spikes (and occupancies) occurred at times before **outcome** (left) or during **reward consumption** (right) to further show that these cells are selective to trial events. **G) Example CA1 neurons in physical space and behavioural task.** Cell 1 fired at the bottom initiation port in Layout Type 2 but not when this same port acted as a B choice in Layout Type 3 or when the port was not a part of the current problem but was visited in Layout 1. Cell 2 fired at one of the B ports in one Layout Type, had no selectivity to the same port in Layout Type 3 when this port was an initiation port and instead fired at a different B choice in Layout Type 3.

During recording sessions (7 to 16 sessions per mouse; 341 to 650 trials per session), we used ten different port layouts, but to simplify the analysis they were all reflections of three basic layout types (Figure 2B), each of which occurred once in every session. In the first layout type, the initiation port (I1) was the top or bottom port, and the choice ports were the far left and far right ports. One of these choice ports remained in the same location in all three layouts used in a session, and will be referred to as the A choice. This acted as a control for physical location, allowing us to assess how the changing context of the different problems affected the representation of choosing the same physical port. Both the other choice port (B choice), and the initiation port, moved physical locations between problems. In the second layout type, both the initiation port (I2) and B choice port (B2) were in locations that were not used in layout type 1. In the third layout type, the initiation port was the same as the initiation port in layout type 1 (I3 = I1), and the B choice port was the same as the initiation port from layout type 2 (B3 = I2). Hence, in every recording session, we had examples of (1) the same port playing the same role across problems, (2) different ports playing the same role across problems and (3) the same port playing different roles across problems (I3 and B2). In addition to these task design controls, we further ruled out the possibility that either potentially invariant aspects of task execution (e.g., velocity and acceleration) or problem/trial specific features (animals’ precise nose position) might explain differences in representations between different brain areas by tracking animal location frame-by-frame using pose estimation software and using linear modelling to account for these variables (Supplementary Figure 7; *Additional Controls for Physical Movement Methods*). The order of the layout types was randomised in each recording session.

As animals transferred knowledge of the trial structure across problems, we reasoned that neurons might exhibit ‘problem-general’ representations of the abstract stages of the trial (initiate, choose, outcome) divorced from the sensorimotor specifics of each problem. On inspection, such cells were common in PFC (Figure 2D). To respond flexibly when a novel problem with the same trial structure is encountered, abstract knowledge should be mapped onto the sensorimotor specifics of the new experience. In line with this, although we observed some problem-general firing in CA1, hippocampal cells were more likely to respond to the specifics of each problem (Figure 2E). We further show that neither PFC nor CA1 cells were selective to physical space *per se* (‘place cell’ like firing fields) but instead were selective to relevant ports and trial events (Figure 2F, G - for more single unit examples see Supplementary Figure 6A, B).

These single unit examples suggest that although problem general representations might exist in both regions, PFC activity appears to generalise more across problems, while CA1 represents physical location more strongly, and additionally exhibits ‘remapping’ between problems in which neurons change their tuning to both physical location and trial events.

### PFC population trial stage related activity generalises more strongly across problems than CA1

To assess whether our single unit observations hold up at the population level, we sought to characterise how neural activity in each region represented trial events, and how these representations generalised across problems. All our consequent population findings were extremely robust and survived the animal random effects test (see *Statistical Significance Methods*, Supplementary Figure 5).

We first assessed the influence of different trial variables in each region using linear regression to predict spiking activity of each neuron, at each time point across the trial, as a function of the choice, outcome, and outcome x choice interaction on that trial (Figure 3A). As the task was self-paced, we aligned activity across trials by warping the time period between initiation and choice to match the median interval (for more details see *Time Warping Methods* and Supplementary Figure 4). We then quantified how strongly each variable affected population activity as the population coefficient of partial determination (i.e., the fraction of variance uniquely explained by each regressor) at every time point across the trial (Figure 3B). This analysis was run separately for each problem in the session and the results were averaged across problems and sessions. Both regions represented current choice, outcome, and choice x outcome interaction, but there was regional specificity in how strongly each variable was represented. Choice (A vs B) representation was more pronounced in CA1 than PFC (peak variance explained - CA1: 8.4%, PFC: 4.8%, p < .001), whereas outcome (reward vs no reward) coding was stronger in PFC (peak variance explained – CA1: 7.1%, PFC: 12.9%, *p* <.001). Furthermore, choice x outcome interaction explained more variance in CA1 than PFC (peak variance explained – CA1: 3.7%, PFC: 2.4%, *p* <.001).

**Figure 3:**
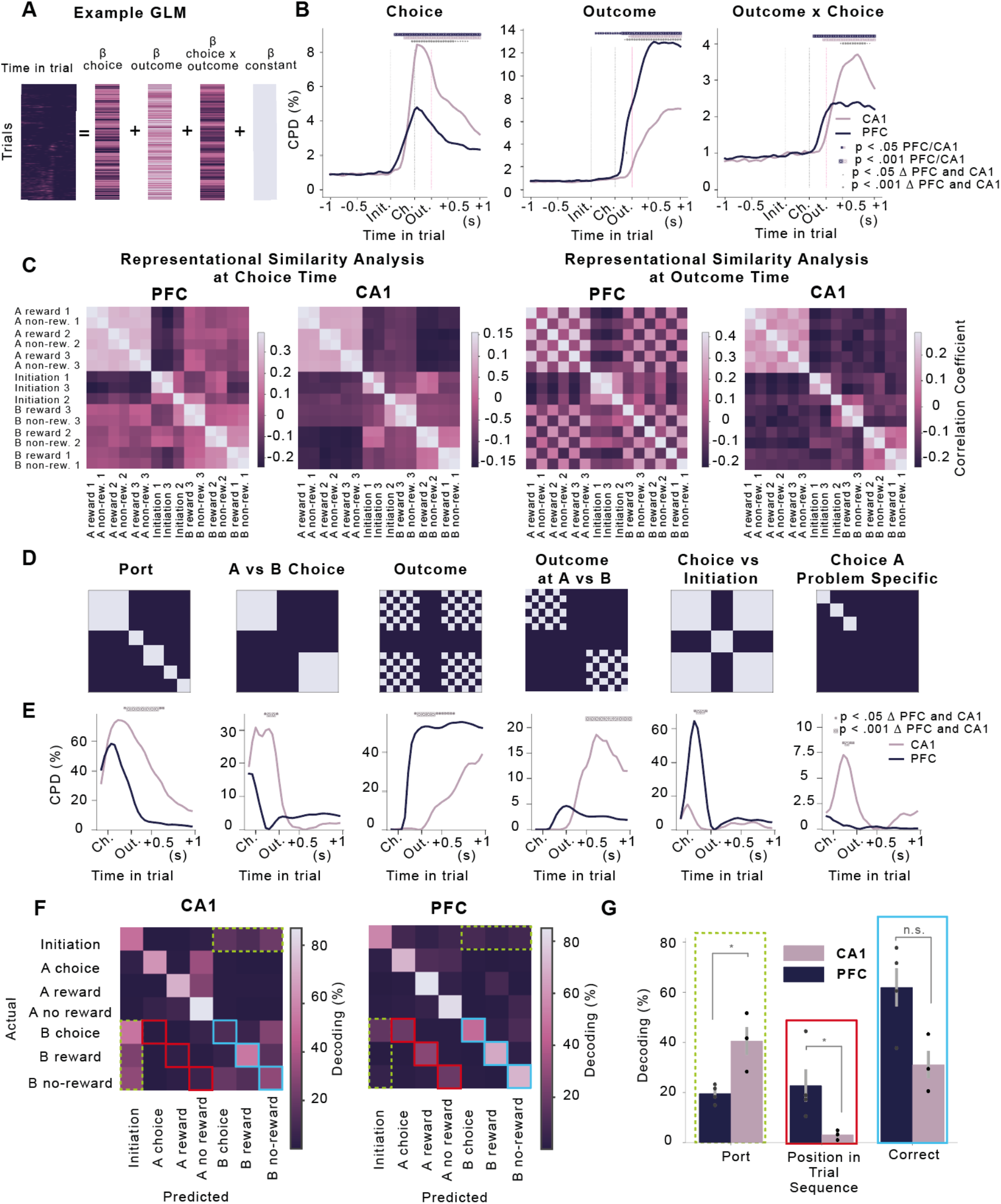
Problem-general and problem-specific representations in PFC and CA1 population activity. **A)** Linear regression predicting activity of each neuron at each time point across the trial, as a function of the choice, outcome and outcome x choice interaction. **B)** Coefficients of partial determination from the linear model shown in **A** for choice, outcome and outcome x choice regressors in PFC and CA1. Significance levels for within region effects were based on a permutation test where firing rates were rolled with respect to trials. Significance levels for differences between regions were based permutation test across sessions corrected for multiple comparison over time points. **C)** Representation similarity at ‘choice time’ (left) and ‘outcome time’ (right), quantified as the Pearson correlation between the demeaned neural activity vectors for each pair of conditions. **D)** Representational Similarity Design Matrices (RDMs) used to model the patterns of representation similarity observed in the data. Each RDM codes the expected pattern of similarities among categories in **C** under the assumption that the population represents a given variable. The *Port* RDM models a representation of the physical port poked (e.g., far left) irrespective of its meaning in the trial. *A vs B Choice* models a representation of A/B choices irrespective of physical port. The *Outcome* RDM models representation of reward vs reward omission. The *Outcome at A vs B* RDM models separate representations of reward vs omission following A and B choices. *Choice vs Initiation* models representation of the stage in the trial. *Choice A Problem Specific* models separate representation of the A choice in different problems. **E)** Coefficients of partial determination in a regression analysis modelling the pattern of representation similarities using the RDMs shown in **D**. The time-course is given by sliding the windows associated with choices from being centred on choice port entry to 0.76 s after choice port entry, while holding time windows centred on trial initiations fixed. Stars indicated time points where regression weight for each RDM was significantly different between the two regions (*p* <.05 (small stars) and *p* <.001 (big stars), permutation test across sessions corrected for multiple comparison over time points. **F)** Confusion matrices from linear decoding of position in trial, using a decoder that was trained on one problem and tested on another, averaged across animals and across all problem pairs. Coloured squares indicate three possible patterns of decoding that indicate different neuronal content. Blue indicates correct cross-task decoding to the same abstract state (e.g., B choice decodes to B choice). Red indicates decoding to a different state that could have occurred at the same sequential position in the trial (e.g., B choice decodes to A choice). Dashed green reflects decoding to the same physical port (e.g., B choice decodes to initiation for task pairs when the B and initiation ports have interchanged between the training and test data, i.e., Layout 2 ➔ Layout 3 and Layout 3 ➔ Layout 2). **G)** Bar plots showing the probability of the cross-task decoder outputting the correct abstract state (blue), the state that has the same physical port as the training data (green - computed only from confusion matrices where B choice and initiation ports interchange), or the other state that can have the same position in the trial sequence (red), computed using the corresponding cells highlighted in **F**. Error bars report the mean ± SEM across different mice. Significance levels were compared against the null distribution obtained by shuffling animal identities between regions.

Though highlighting some differences in population coding between regions, this approach cannot assess the relative contribution of abstract representations that generalise across problems versus features specific to each problem, such as the physical port location. This requires comparing activity both across time points in the trial and across problems, which we did using representational similarity analysis (RSA)^44^. We extracted firing rates around initiation and choice port entries (+/- 20ms around each port entry type) and categorised these windows by which problem they came from, whether they were initiation or choice, and - for choice port entries whether the choice was A or B and whether it was rewarded - yielding a total of 15 categories (Figure 3C). For each session we computed the average activity vector for each category, then quantified the similarity between categories as the correlation between the corresponding activity vectors. We show RSA matrices for this ‘choice time’ analysis (Figure 3C, left panels), and an ‘outcome time’ analysis (Figure 3C, right panels) where the windows for choice events were moved 250ms after port entry, holding the time window around trial initiations constant.

To quantify the factors influencing representation similarity, we created representational similarity design matrices (RDMs) which each encapsulated the predicted pattern of similarities under the assumption that activity was influenced by a single trial/problem feature (Figure 3D). For example, if the population activity represented only which physical port the animal was at, its correlation matrix would look like Figure 3D, Port. We included design matrices for a set of problem-general features; the trial stage (‘Initiation vs Choice’), choice (A vs B), trial outcome (both on its own as ‘Outcome’, and in conjunction with choice ‘Outcome at A vs B’). Changes in activity across problems might occur simply due to neurons being tuned for particular physical port locations, which will be captured by the ‘Port’ RDM. However, it is also possible that the changing context provided by different problems modifies the representation of choosing the same physical port at the same trial stage. To assess such ‘remapping’, we included an RDM ‘Choice A problem specific’ which modelled problem specific representations of the A choice, which shares the same physical port location and meaning across problems. We modelled the observed pattern of similarities in the data as a linear combination of these RDMs, quantifying the influence of each by its corresponding weight in the linear fit. To be able to examine the temporal evolution of these effects we ran a series of regressions onto the data. In each, the data around initiation port entry was the same but the data around the choice port entry progressed serially through time from choice point until after the reward was delivered (Figure 3E).

Consistent with our single unit observations, both PFC and CA1 represented both problem-specific and problem-general features to some extent. However, there was a marked regional specificity in how strongly different features were encoded (Figure 3E). PFC had stronger, abstract, sensorimotor-invariant representation of trial stage (initiation vs choice) and trial outcome (*p* <.001). In contrast, CA1 had stronger representation of the physical port the subjects were poking, and whether it was an A vs B choice (*p* <.001). Additionally, CA1 but not PFC showed a problem specific representation of A choices (*p* <.001). This is striking because during A choices both the physical port and its meaning are identical across problems, indicating that the changing problem context alone induced some ‘remapping’ in CA1 but not PFC. Finally, there was a regional difference in the representation of trial outcome. PFC outcome representations were more general (the same neurons responded to reward or reward omission across ports and problems – *p* <.001). CA1 also maintained an outcome representation, but this was more likely to be conjunctive than in PFC – different neurons would respond to reward on A and B choices (*p* <.001). To ensure that these regional differences in representation were not driven by fine-grained selectivity to physical movements, we re-ran the analysis on residual firing rates after the effect of velocity, acceleration and 2D nose position were regressed out (for more details see *Additional Controls for Physical Movement Methods*). All inter-region differences except the stronger representation of A vs B choice in CA1 survive this control (Supplementary Figure 7), consistent with the single cell examples described above (Figure 2G, F; Supplementary Figure 6). We also assessed whether problem-specificity in CA1 might be driven by slow drift over time, but found that representations changed abruptly at transitions between problems (Supplementary Figure 11).

To further characterise differences in representation between regions, we trained a linear model to decode different positions in the trial (Initiation, A choice, A reward, A no-reward, B choice, B reward and B no-reward) on one problem, and tested the decoding performance on a different problem. Because the B and initiation ports moved between problems, and sometimes interchanged, we were able to dissociate port-specific and abstract task state decoding patterns. PFC predominantly decoded to the correct task state (Figure 3F-G). Where PFC made errors, they were predominantly to the other state that could occur at the same sequential position in the trial (A rather than B choice or outcome). By contrast, CA1 predominantly decoded to the same physical port as the training data. Together, these population results confirm that PFC had a predominantly generalising representation, and this representation embeds the sequential properties of the trial while CA1 encoded problem specifics (such as port identity) more strongly.

### Low dimensional temporal structure of activity is invariant across problems and regions, but cell assemblies generalise more strongly in PFC than CA1

To further explore how the structure of population activity generalised between problems, we assessed how accurately low dimensional activity patterns in one problem could explain activity in another. Using singular value decomposition (SVD), we decomposed activity in each problem into a set of cellular and temporal modes. Cellular modes correspond to sets of neurons whose activity covaries over time, and hence can be thought of as cell assemblies. Each cellular mode is specified by a vector with a weight for each cell, indicating how strongly the cell participates in the mode. Cellular and temporal modes come in pairs, such that each cellular mode has a corresponding temporal mode, which is a vector of weights across time-points indicating how the activity of the cellular mode varies over time.

To evaluate the cellular and temporal modes for a given problem, we first regressed out general movement-related features onto the firing rates (for more details see Supplementary Figure 7, *Additional Controls for Physical Movement Methods*). After removing the effect of velocity, acceleration and 2D nose position we computed the average residual firing rate at each time point across the trial, for four types of trials – rewarded A choices, non-rewarded A, rewarded B, and non-rewarded B (non-rewarded trials included both correct trials and incorrect trials). For each cell, we concatenated these 4 time series to create a single time series containing the average activity of the cell across each time point of the four trial types. The temporal modes span this same set of time points and hence capture variation across both timein-trial and trial-type. We then stacked these single cell activity time series for all neurons, to create an activity matrix *D* where each row contained the activity of one neuron (Figure 4A). Using SVD, we decomposed this activity matrix into cellular and temporal modes *U* and *V*, linked by a diagonal weight matrix ∑

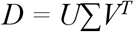

**Figure 4:**
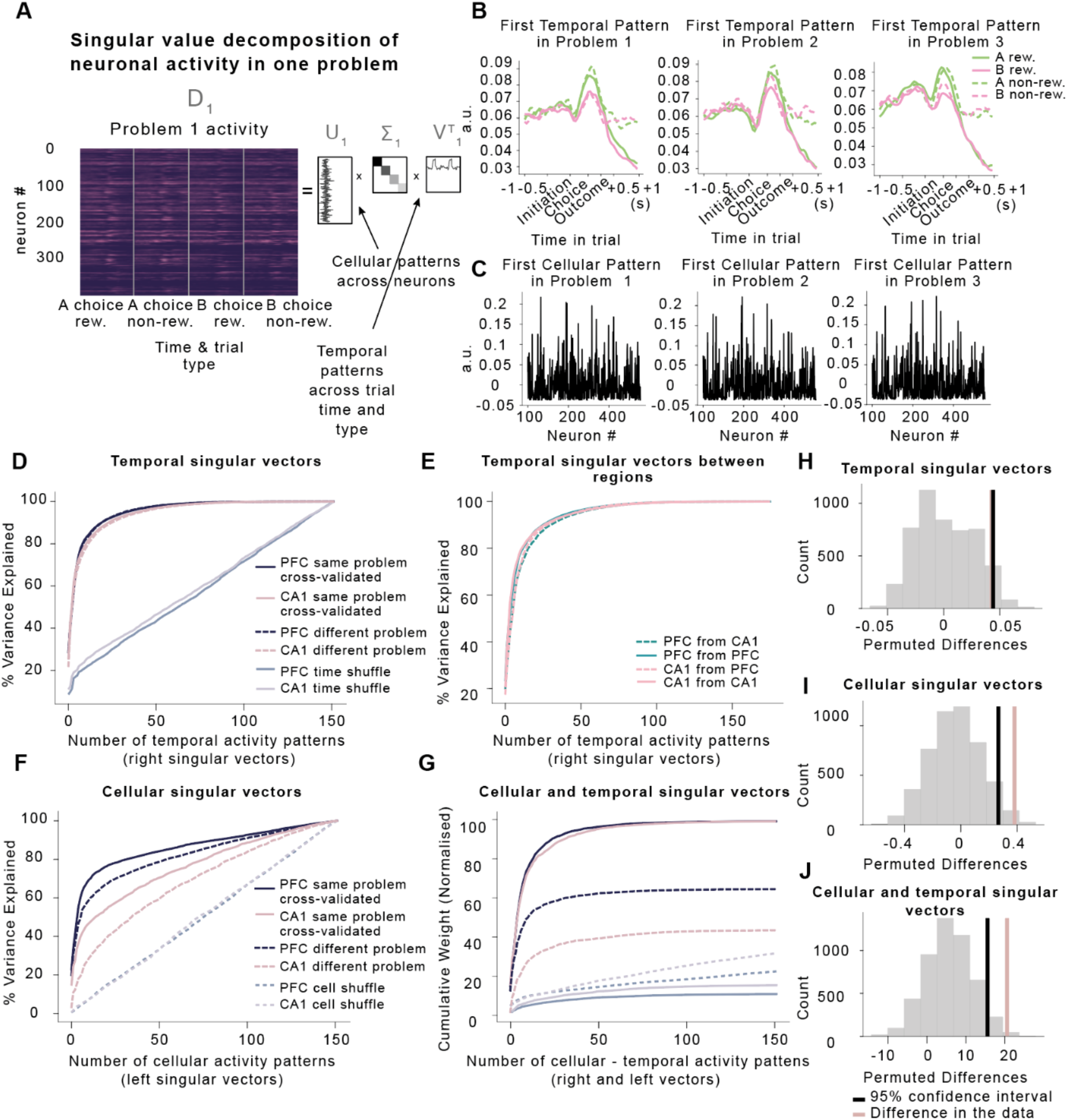
Generalisation of low dimensional representations of trial events. **A)** Diagram of singular value decomposition (SVD) analysis. A data matrix comprising the average activity of each neuron across time points and trial types was decomposed into the product of three matrices, where diagonal matrix ∑ linked a set of temporal patterns across trial type and time (rows of *V^T^*) to a set of cellular patterns across cells (columns of *U*). **B)** First temporal mode in *V^T^* from SVD decomposition of data matrix from PFC plotted in each problem separately for clarity and separated by A (green) and B (pink) rewarded (solid) non-rewarded (dashed) choices. **C)** First cellular mode from SVD decomposition of data matrix from PFC in each problem showing similar pattern of cells participate in all problems. **D)** Variance explained when using temporal activity patterns *V_1^T^_* from one problem to predict either held out activity from the same problem (solid lines) or activity from a different problem (dash lines). Light purple and lilac lines indicate variance explained when shuffling time points in the firing rates matrices. **E)** Variance explained when using temporal activity patterns *V_1^T^_* to predict either activity from the same problem and brain region (solid lines) or a different brain region (and therefore different animal) and the same problem (dash lines) *D_2_*. **F)** Variance explained when using cellular activity patterns *U_1_* from one problem to predict either held out activity from the same problem (solid lines) or activity from a different problem (dash lines). Dashed light purple and lilac lines indicate variance explained when shuffling cells in the firing rates matrices. **G)** Cumulative weights along the diagonal ∑ using pairs of temporal *V_1^T^_* and cellular *U_1_* activity patterns from one problem to predict either held out activity from the same problem (solid lines) or activity from a different problem (dash lines). Weights were normalised by peak cross-validated cumulative weight computed on the activity from the same problem. **H)** To assess whether the temporal singular vectors generalised significantly better between problems in PFC than CA1, we evaluated the area between the dash and solid lines in **D** for CA1 and for PFC separately, giving a measure for each region of how well the singular vectors generalised. We computed the difference in this measure between CA1 and PFC (pink line in **H**), and compared this difference to the null distribution obtained by permuting sessions between brain regions (grey histogram, black line shows 95th percentile of distribution). Temporal singular vectors generalised equally well between problems in the two regions. **I)** Cellular singular vectors generalised significantly better between problems in PFC than CA1. Computed as in **H** but using the solid / dash lines from **F**. **G)** Pairs of cellular and temporal singular vectors generalised significantly better between problems in PFC than CA1. Computed as in **H** but using the solid / dash lines from **G**.

The cellular modes are the columns of *U*, and the temporal modes are the rows of *V^T^*. Both modes are unit vectors, so the contribution of each pair to the total data variance is determined by the corresponding element of the diagonal matrix ∑. The modes are sorted in order of explained variance, such that the first cellular-temporal mode pair explains the most variance. The first cellular and temporal mode of PFC activity in three different problems is shown in Figure 4B, C. It is high throughout the ITI and trial with a peak at choice time, but strongly suppressed following reward (similar to cell 5 in Figure 2D).

We reasoned that: (i) if the same events were represented across problems (e.g. initiation, A/B choice, outcome), then the temporal modes would be exchangeable between problems, no matter whether these representations were found in the same cells; (ii) if the same cell assemblies were used across problems, then the cellular modes would be exchangeable across problems, no matter whether the cell assemblies played the same role in each problem; and (iii) if the same cell assemblies performed the same roles in each problem, then pairs of cellular and temporal modes would be exchangeable across problems.

To see whether the same representations existed in each problem, we first asked how well the temporal modes from one problem could be used to explain activity from other problems. Since the set of temporal modes *V* is an orthonormal basis, any data of the same rank or less can be perfectly explained when using all the temporal modes. However, population activity in each problem is low dimensional so a small number of modes explain a great majority of the variance. Modes that explain a lot of variance in one problem will only explain a lot of variance in the other problem if the structure captured by the mode is prominent in both problems. The question is therefore how quickly variance is explained in problem 2’s data, when ordering the modes according to variance explained in problem 1. To assess this, we projected the data matrix *D_2_* from problem 2 onto the temporal modes *V_1_* from problem 1, giving a matrix *M_V_* whose elements indicate how strongly each temporal mode contributes to the problem 2 activity of each neuron:

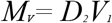

The total variance explained by each temporal mode is given by squaring the elements of *M_V_* and summing over neurons. We plot the cumulative variance explained as a function of the number of temporal modes used (Figure 4D). To control for drift in neuronal representations across time, we computed the data matrices separately for the first and second halves of each problem. We compared the amount of variance explained using modes from the first half of one problem to model activity in the second half of the same problem, with the variance explained using modes from the second half of one problem to model activity from the first half of the next problem.

In both PFC and CA1, the cumulative variance explained as a function of the number of temporal modes used, did not depend on whether the two data sets were from the same problem (solid) or different problems (dashed) (Figure 4D, H, *p* >.05). This indicates that the temporal patterns of activity, and therefore the trial events represented, did not differ across problems in either brain area. However, as this analysis used only the temporal modes, it says nothing about whether the same or different neurons represented a given event across problems. In fact, we can even explain activity in one brain region using temporal modes from another region and mouse (Figure 4E).

The pattern was very different when we used cellular modes (i.e., assemblies of co-activating neurons) from one problem to explain activity in another. We quantified variance explained in problem 2 using cellular modes from problem 1, by projecting the problem 2 data matrix *D_2_* onto problem 1 cellular modes *U*_1_, giving a matrix *M_u_* whose elements indicate how strongly each cellular mode contributes to problem 2 the activity at each timepoint:

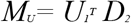

The total variance explained by each temporal mode is given by squaring the elements of *M_U_* and summing over timepoints. In both PFC and CA1, cellular modes in *U* that explained a lot of variance in one problem, explained more variance in the other half of the same problem than they did in an adjacent problem (Figure 4F - differences between solid and dashed lines). However, the within problem vs cross problem difference was larger in CA1 than PFC (Figure 4I, *p* <.05). This indicates that PFC neurons whose activity covaried in one problem were more likely to also covary in another problem, when compared to CA1 neurons. As this analysis considered only the cellular modes it does not indicate whether a given cell assembly carried the same information across problems.

To assess how well the cellular-temporal activity patterns from problem 1 explained activity in problem 2, we projected data set *D*_2_. onto the cellular and temporal mode pairs of problem 1 (*U_1^T^_, V_1_*).

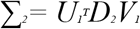

If the same cell assemblies perform the same roles in two different problems, the temporal and cellular modes will align, and ∑_2_ will have high weights on the diagonal. We therefore plotted the cumulative squared weights of the diagonal elements of ∑ within and between problems (Figure 4G). In both PFC and CA1 cellular and temporal modes aligned better in different data sets from the same problem (solid lines), than for different problems (dashed lines). However, this difference was substantially larger for CA1 than PFC (Figure 4J, *p* <.05). All results also held true when using a time window between only initiation and choice (Supplementary Figure 8).

These data show that although the temporal structure of activity in both regions generalises perfectly across problems, brain regions and subjects – a consequence of the same set of trial events being represented in each, the cell assemblies used to represent them generalised more strongly in PFC than CA1.

### Policy representations are abstract in PFC, but linked to sensorimotor experience in CA1

So far, we have focused on the neuronal representations of events on individual trials, and how they generalise across problems. But to maximise reward, the subject must also track which option is currently best by integrating the history of choices and outcomes across trials. To be useful for generalisation, this policy representation should also be divorced from the current sensorimotor experience of any specific problem.

To obtain an estimate of subjects’ beliefs about which option was best, we used a logistic regression predicting current choices as a function of the history of previous rewards, choices and their interactions (Figure 5A). This allowed us to compute, trial-by-trial, the probability that the animal would choose A vs B – i.e., the animal’s policy. When we used this policy as a predictor of neural activity, it explained variance that was not captured by within-trial regressors such as choice, reward and choice x reward interaction. Specifically, the subjects’ policy interacted with the current choice explained variance (Figure 5B, *p* <.001). Notably, this signal became prominent around the time of trial initiation, when it would be particularly useful for guiding the decision.

**Figure 5:**
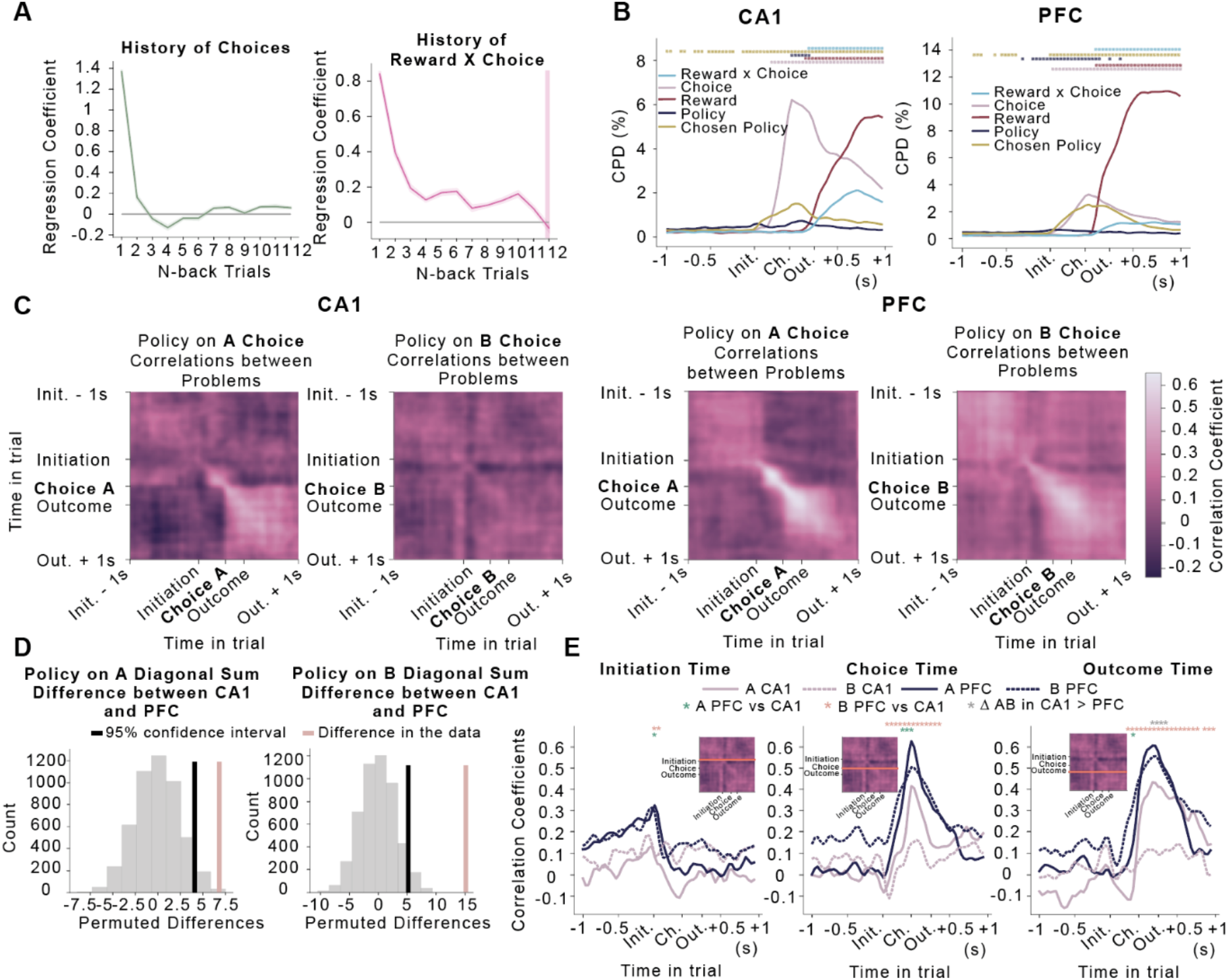
Policy Generalisation in PFC and CA1. **A)** Weights from logistic regression predicting choices in recording sessions using choices, rewards and choice x reward interactions over the previous twelve trials as predictors. The effect of choice x outcome interaction history was significantly above zero on up to eleven trials back (one sampled t-test, *p* < .05) except for the 7^th^ trial (*t* _(6)_ = 1.86, *p* = .112). Error bars report the mean ± SEM across mice. **B)** CPDs from regression models predicting neural activity using current trial events, subjects’ policy (estimated using the behavioural regression in **A**, and policy interacted with current choice. Stars denote the time points at which each regressor explained significantly more variance than expected by chance (permutation test based on rolling firing rates with respect to trials – *p* <.001, corrected for multiple comparisons; for more details on permutation tests see *Statistical Significance Methods*.) **C)** Correlations across problems between policy weights in regressions predicting neural activity. Regressions were run separately for A (left panels) and B (right panels) choices in each problem, and at each time point across the trial. Correlations of policy representations between all problem pairs were evaluated for each pair of time points, values on the diagonal show how correlated policy representation was at the same time point in both problems. Positive correlation indicates that the same neurons coded policy with the same sign in both problems. **D)** To quantify whether policy generalised more strongly between problems in PFC than CA1, we computed the between region difference in the sum along the diagonal of the correlation matrices in **C)**, separately for A and B choices, and compared it against the null distribution obtained by permuting sessions between brain regions. Policy representation on both A and B choices generalised more strongly in PFC than CA1. **E)** Slices through the correlation matrices at initiation (left), choice (centre) and outcome (right) times for A (solid) and B (dash line) choices. Significant differences between conditions are indicated by stars as shown in legend.

To examine whether policy representations generalised across problems, we evaluated the correlation across problems between the policy weights in the neural regression. Because the A port was the same on each problem, but the B port varied between problems, we computed policy regression weights at each time point separately for A and B choices (controlling for reward). We then computed the average across-problem correlation of these weights between every pair of timepoints (Figure 5C). The diagonal elements of these matrices show the average correlation across problems at the same time point in each problem. Visual inspection (Figure 5C), and permutations tests of differences between sums of the diagonals of Policy on A and B choices correlation matrices (*p* <.05), revealed that these correlations were larger in PFC than CA1 (Figure 5D). On average, therefore, cellular representations of policy generalised across problems better in PFC than CA1 on both A and B choices.

One possible explanation is that PFC simply represented action values in a problem-general way. A more interesting possibility is that current policy shapes the representation of each trial stage differently, but in CA1 these representations are more tied to the sensorimotor specifics of the current problem. To test this, we examined time-slices through the correlation matrices at initiation, choice, and outcome times (Figure 5E). In PFC, all three correlation profiles on both A and B trials peaked at the correct time point (the equivalent to the diagonal elements of the matrix) – i.e., the policy representations generalised across problems, but were specific to the different parts of the trial (initiate, choose, outcome). A similar pattern was present in CA1, but only on A choices (which are the same physical port across problems). No CA1 correlation was significantly above zero on B choices. Indeed, whilst PFC policy correlations were greater than CA1 correlations for all representations (all *p* <.05) on both A and B choices, CA1 correlations showed a greater difference between A and B trials at outcome time (Figure 5E, all *p* <.05).

Overall, therefore, both PFC and CA1 maintained representations of the subject’s current policy that were not simple value representations – as they differed depending on the trial stage. These representations were abstracted across problems in PFC, but tied to the sensorimotor specifics in CA1. A portion, but not all, of this problem specificity in CA1 was accounted for by the port identity.

## 3 Discussion

Humans and other animals effortlessly generalise prior experience to novel situations that are only partially related. To do this, we must reduce experiences to abstractions - features that are common between different situations. Critically, we must also bind these temporally extended abstractions to the incoming sensory particularities of the current situation. Our study makes three different types of contribution to understanding how, and when, this process happens.

First, we show that this focus on abstraction, common in studies of spatial reasoning and memory^2,3^ is also important in standard reinforcement learning paradigms such as reversal learning. Where the dominant focus in these paradigms has been on variables such as value and prediction error^45-47^ (important for learning actions *de novo*), we show that the neural representation in mPFC reflects the temporal structure of the problem itself (spatial or non-spatial), that may allow actions to be generalised from previous similar experiences. One intriguing possibility is that such representations are formed during the shaping process that precedes most operant experiments.

Second, we show that mPFC and CA1 contain different representations that suggest different functional roles. Population responses in mPFC were dominated by problem-invariant representations that might form the abstraction. By contrast, the CA1 responses contained major sources of variance that were either invariant to the sensorimotor particularities (port selective), or intriguingly, the interaction of these with the problem structure (demonstrating ‘remapping’ between problems, or reflecting the interaction of task policy and individual port). Representations such as these are required to bind task-general abstractions to the current sensory problem.

Third, we show that task abstractions in mPFC simultaneously represent behaviour over markedly different temporal scales. Part of the mPFC representation pertained to the immediate next action in the sequence (e.g., go to the initiation port), but part of the representation pertained to the integrated history of rewards and actions over many trials, that allowed the animal to make profitable choices. Notably, both parts of the representation were maintained in an abstract form (albeit with a small port component) that generalised over problems with different sensory particularities.

These findings are related to previous findings across a number of different literatures.

In reinforcement learning, recent data have highlighted the low dimensional structure of abstract task representations in rodent OFC^9^. This aligns with our finding that low dimensional ***temporal modes*** are consistent across different sensorimotor instances of the reversal learning problem in both mPFC and CA1. We also confirm that they are consistent between animals and further demonstrate they are broadly consistent between different brain areas (mPFC and CA1), suggesting this low dimensional ***temporal*** structure does not reflect the unique representational properties of a particular brain area. Notably, however, because we record across the same neurons in different problems, we are able to ask not only whether the temporal dimensions are preserved across problems, but whether these temporal modes align to the same neurons in each problem, i.e., whether the same neurons represent the same trial events across problems. They do so significantly more in PFC than CA1. It is this that enables us to propose different functional roles for the two different brain regions.

Recent RL work has also found a form of abstraction in primate PFC and hippocampus^48^. Because abstraction was assessed across conditions that used the same physical operandum and hence shared sensorimotor correlates, it is not possible in these data to discern whether hippocampal representation would generalise to different sensorimotor instantiations of the same problem. By contrast, the focus of our study is on how these brain regions enable generalisation of knowledge across problems the ***same abstract structure*** but ***different*** sensorimotor experiences.

The essence of RL is the integration of rewards over temporally extended experiences to generate expected values or policies^49^. Our demonstration that these policy representations are abstracted aligns directly with ideas from computer science such as meta-reinforcement learning^6,50,51^, which have recently been proposed as models to understand prefrontal activity. Indeed, our behavioural data directly demonstrate meta-learning, as reversals become faster with increasing experience.

Notably, we also found that policy coding was not unique to prefrontal cortex, as hippocampus also contained policy representations, corroborating existing findings for the existence of signals relevant for decision-making in hippocampal formation^52,53,54^. We expand on these observations to provide further evidence that hippocampal activity might represent sensorimotor specifics of events in the context of broader memory schemas and task structures.

Whilst relatively new to the neuroscience of RL, the overarching ideas in our study are central in the study of memory and space. Here it is commonly assumed that hippocampal representations reflect the sensory details of each episodic experience^19,20,55^, and cortical representations abstract these details to allow generalisation ^56,57,58^. Indeed, in spatial studies in rodents, new abstractions (schemata) rely causally on mPFC^3^. Equally, spatial reasoning in rodents is dependent on grid cells^30^, which abstract of the fundamental 2D properties of physical space. Recent data and modelling have shown that hippocampal spatial representations are bound to this abstraction^59,40^. We believe that this study demonstrates that many of these ideas carry directly over to structural abstractions in RL problems, and therefore further align these historically distinct fields.

We do not perceive the world as it really is. Starting with the visual 2D inputs on the retina that we use along with prior experience to infer the 3D world around us^60^, our brains likely develop structural placeholders for many of our experiences. In fact, we remember things more easily if we know the general schema or a script for a particular even^61^, and often ignore information that does not align with our understanding of the world^62^. More broadly, here we demonstrate that mice also acquire sophisticated models of tasks they frequently experience in their environment and can apply this knowledge to solve new problems faster. We further show that prefrontal cortex contains representations of what can be thought of as a ‘learning-set’, or ‘schema’ of abstract relationships and variables needed to solve new related problems while hippocampus combines sensorimotor and abstract information to represent an interaction between the two, which might be crucial for both interpreting our ongoing experiences as well as encoding and recall of episodic memories.

## Author Contributions

V.S., T.A., M.E.W. and T.E.J.B. designed the study; V.S., T.A. and J.L.B. acquired the data; V.S. and T.E.J.B analysed the data with input from T.A. V.S., T.A. and T.E.J.B wrote and edited the manuscript with input from M.E.W.

## Acknowledgments

We would like to thank Tom Jahans-Price for his help with setting up electrophysiology in our lab and training us to conduct our first recordings. We would also like to thank Tom Jahans-Price, Mohamady El-Gaby and Yves Weissenberger for providing helpful comments on the drafts of the manuscript. This work was funded by the following grants: Wellcome Principal Research Fellowship (219525/Z/19/Z), and JS McDonnell Foundation award (JSMF220020372) to T.E.J.B.; Wellcome Collaborator award (214314/Z/18/Z) to T.E.J.B., T. A., M.E.W. and Senior Research Fellowship (202831/Z/16/Z) to M.E.W. The Wellcome Centre for Integrative Neuroimaging and Wellcome Centre for Human Neuroimaging are each supported by core funding from the Wellcome Trust (203139/Z/16/Z, 203147/Z/16/Z).

## Data and materials availability

All data, analysis and behavioural training code will be released on publication.

## 4 Materials and Methods

### Behavioural Apparatus

Experiments were performed in custom made operant boxes, controlled using pyControl^63^ (https://github.com/pyControl/hardware). The boxes used in the training phase of the experiment had six nose poke ports mounted on the back wall, each with infrared beam, stimulus LED and solenoid valve for dispensing liquid rewards, and a speaker for auditory stimuli. For recording experiments mice were transferred to operant boxes with nine nose poke ports located in electrically shielded sound attenuating chambers.

### Subjects

Nine male C57BL/6J experimentally naïve mice bred in the Biomedical Sciences Facility at the University of Oxford were obtained for this experiment at six weeks of age. Animals were group housed prior to surgery, and individually housed post-surgery, in a humidity- and temperature-controlled vivarium, on a 12-hour light-dark cycle (7:00 to 19:00). All nine animals were implanted with silicon probes, but we only obtained data from seven animals, due to one probe being damaged during surgery and having to cull one animal prior to recordings. Experiments were carried out in accordance with the Oxford University animal use guidelines and performed under UK Home Office Project Licence P6F11BC25.

### Behavioural Training

Mice were placed on water restriction 48 hours prior to starting behavioural training, with 1 hour water access provided 24 hours before the first session. Mice were trained six days per week, and on the day off they received 1 hour ad lib water access in their home cage. On training days, mice typically received all their water in the task, but were given additional water if required to maintain their body weight above 85% of their pre-restriction baseline weight.

Mice were trained on a sequence of reversal learning problems each with the same structure but a different physical port layout. Each reversal learning problem used three nose poke ports, out of the six or nine ports available in the operant box. One port was used for trial initiation, the other two were choice ports where reward could be obtained. During the initial training phase (Figure 1A) ports not used in the current problem were covered. During recording sessions, ports used in all three problems presented in the session were exposed throughout, and unused ports were covered.

Each trial started with the initiation port lighting up, until the subject poked it, after which two choice ports both lit up. Mice chose one of the choice ports which triggered a sound cue (250ms long) indicating the trial outcome, with a pure tone (5 kHz) indicating they will get a reward and white noise indicating reward omission. Reward was delivered at the termination of the auditory cue. A 2s inter-trial interval started once the animal left the port following reward consumption or a non-rewarded choice. One in four randomly selected trials was a forced choice trial, where a single randomly selected choice port lit up which the animals had to select. At any given point in time, one choice port had a high reward probability and the other one had low probability. Reward probability reversals were triggered 5-15 trials after the subject crossed a threshold of 75% correct choices (exponential moving average, tau=8 trials).

In the initial training stage of experiment, mice (Figure 1) encountered a single problem (i.e., port layout) per session, and moved to the next problem the session after they had completed 10 reversals on the current problem. In each problem, the first three reversals had reward probabilities of 0.9 and 0.1 at the good/bad choice ports. The fourth and fifth reversals had reward probabilities of 0.85 and 0.15, and the remaining reversals had reward probabilities of 0.8 and 0.2. In this phase each session was 30 minutes long and animals performed two sessions per day. The reward sizes during this stage were incrementally decreased from 15 ul in the beginning of the training to 4 ul, based on the current weight of the animal and its performance on the previous session. Each session started with a free reward given from each of the two choice ports. Mice were divided into three groups with each group starting on a different layout. Sequentially presented layouts were chosen to be as different as possible, and the sequence of problem layouts was counterbalanced across animals.

Once mice had completed 10 problems during this initial training phase, we started to present multiple problems in each session, to prepare them for recording sessions where we sought to record neurons across multiple problems. Initially, mice were trained on two problems in a session, in the nine port operant boxes subsequently used for recordings. Mice completed 12 different problems in this stage, with the port layout used in each chosen to be as different from the previous one as possible. The reward probabilities in this phase were always 0.8 and 0.2 and the reward size was 4 ul. After mice completed two reversal blocks on one layout, choice ports that were going to be a part of the new problem layout both lit up. Mice received a free reward from each of the new choice ports. Next, the new initiation port lit up signalling mice where they could initiate a trial. For a schematic of all port layouts and counterbalancing used in all stages of the experiment see Supplementary Figure 9.

### Behavioural Training during Recordings

During recordings, subjects completed four reversal blocks in each of three different problem layouts in every session. All task parameters (including forced trials) were kept the same as during the two-layout per session training stage, with the exception that now subjects needed to complete four blocks on each problem before they were moved onto a new one. As before, the problem layout change was signalled by the two new choice ports lighting up and staying lit up until the subject collected a reward from each port. This was followed by the new initiation port lighting up. Port layouts used during recording sessions were designed to allow us to ask specific questions of the neural activity. As described in the Results section, all layouts were reflections of three basic layout types, each of which was presented once each session, in a randomised order (Figure 2B).

### Electrophysiological Recordings and Spike Sorting

The silicon probes used were Cambridge Neurotech 32 channel probes. F series probes were used for hippocampus, P series for mPFC. For hippocampal recordings we started the recordings only after we lowered the probe enough to detect characteristics of hippocampus sharp wave ripples in the local field potential while the animal was asleep in its home cage. For mPFC recordings we lowered the probe ~100um on every recording day. For more details on recording sites, see Supplementary Figure 3. Neural activity was acquired at 30kHz with a 32-channel Intan RHD 2132 amplifier board (hardware bandpass filtering between 1.1 and 7603.8 Hz; Intan Technologies, USA) connected to an OpenEphys acquisition board via a flexible serial peripheral interface cable (‘Ultra Thin RHD2000 SPI cable’, Intan Technologies). Behavioural and ephys data were synchronised by sending sync pulses from the pyControl system to the OpenEphys acquisition board. Electrophysiological recordings were then spike sorted offline using KiloSort^64^ and manually curated using phy (https://github.com/kwikteam/phy). Clusters were classified as single units and retained for further analysis if they had a characteristic waveform shape, showed a clear refractory period in its autocorrelation, were stable over time and were present only on nearby channels. We merged clusters only if there was a high similarity in waveforms and channels they came from, had a refractory period in their cross-correlation histograms and occupied similar areas in feature space or appeared to drift into one another.

### Surgery and Histology

Subjects were taken off water restriction 48 hours prior to surgery, then anaesthetised with isoflurane (3% induction, 0.5–1% maintenance), treated with buprenorphine (0.1 mg/kg) and meloxicam (5 mg/kg), and placed in a stereotactic frame. A silicon probe mounted on a Microdrive (Ronal Tools) was implanted into either mPFC (AP:1.95, ML:0.4, DV:-0.8), or dCA1 (AP:-2, ML:1.7, DV:-0.7), and a ground screw was implanted above the cerebellum. Both of the DV coordinates are relative to the brain surface. Mice were given additional doses of meloxicam each day for 3 days after surgery, and were monitored carefully for 7 days postsurgery, then placed back on water restriction 24 hours before restarting task behaviour. At the end of the experiment, electrolytic lesions were made under terminal pentobarbital anaesthesia to mark the probe location, animals were perfused, and the brains fixed in formal saline for subsequent histology to identify lesion locations.

### Data analysis

All analyses were carried out using custom written code in Python. Only sessions where animals completed three problems and four reversals in each problem were used for subsequent neural data analyses. This selection criteria ensured that in sessions used for neural data analysis both the position of the port and the passage of time were controlled for.

### Time in Trial Alignment

Activity was aligned across trials by warping the time interval between trial initiation and choice to match the median interval across all recorded trials. Activity prior to trial initiation or after choice was not warped. Spike times that occurred between initiation and choice were converted into the aligned reference frame by linear interpolation between initiation and choice time. The firing rate of each neuron was calculated in the aligned reference frame at time points evenly spaced every 40ms, from 1 second before trial initiation to 1 second after trial outcome, using a Gaussian kernel with 40ms standard deviation. To compensate for the change in spike density due to time warping, spikes in the warped interval between initiation and choice were weighted by the stretch factor applied, prior to evaluating the firing rate. For an illustration of the process details see Supplementary Figure 4.

### Statistical Significance

The significance of the differences between brain areas in analyses reported throughout the paper was computed by shuffling the sessions of CA1 and PFC animals to obtain null distributions. To correct for multiple comparisons, the null distributions were formed by taking the peak difference between CA1 and PFC all time points in each permutation. This approach is a commonly used method for family-wise error correction for permutation tests^65^. Real differences in the data were compared against the 95th and 99th percentiles of such null distributions. All comparisons also survived a group test obtained by shuffling animal identities between regions (Supplementary Figure 5). To establish the significance levels for the effects within regions (Figure 3B, Figure 5B), the firing rates were rolled with respect to trial identities, so that the autocorrelations between consequent trials were retained.

### Representational Similarity Regression Analysis

We created representational similarity matrices which consisted of the Pearson correlation coefficients of neurons in 15 different conditions, defined by the trial stage, choice, outcome, and problem number (see Results section and Figure 3). Because neurons were not simultaneously recorded, we collapsed data across recording sessions for each brain region into a single matrix (cells x trial events) and then calculated the correlation matrix across cells between different trial events (i.e., representational similarity). We used a linear regression to model the patterns of representation similarity in the data as a linear combination of representation similarity design matrices (RDMs):

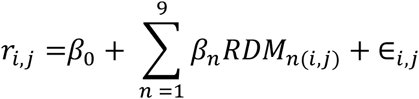

Where r_(i,j)_ are elements of the RSA matrix and *RDM*_n(i,j)_ are elements of the *n*th RDM. The set of RDMs used is shown in Figure 3D. Before regressing the correlation matrices onto the RDMs the diagonal elements from both were deleted and a constant matrix of ones was added to the design matrix to account for any condition independent correlation between neurons. We plotted the coefficients of partial determination (CPDs) from the regression model described above. The CPD was defined as:

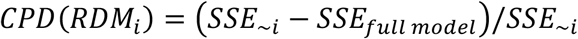

Where *SSE_~i_* refers to the sum of squares from a regression model excluding the RDM_i_ of interest and *SSE*_full model_ is the sum of squares from a regression model including all the RDMs. CPDs describe how much unique variance each RDM accounts for in the RSA matrix calculated from firing rates.

### Surprise Measure

To investigate the time course of how quickly the firing rates of neurons change in response to layout changes (Supplementary Figure 11), we used the ‘surprise’ measure from the information theory:

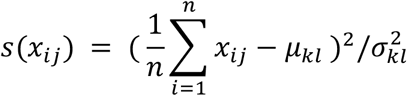

where x_ij_ is the firing rate of one neuron on a given trial *i* and problem layout *j*, *μ* _k_ and *σ* _k_ are the baseline mean and the standard deviation of the firing rate of that neuron on a particular problem layout. If *j* = *k*, then the s(x_ij_) on each trial *i* is calculated based on the mean firing rate *μ* and standard deviation *σ* of the withheld trials from the same problem. More precisely, to calculate how much the firings rates change during the same problem layout s(x_ij_) was calculated on the 10 trials before the problem layout switch (‘test’ within problem), where *μ*_k_ and *σ* _k_ were calculated on the 10 trials before those ‘test’ trials (‘train’ within problem). If *j* ≠ *k*, then the s(x_ij_) on each trial *i* was calculated based on the mean firing rate *μ* and standard deviation *σ* of the withheld trials from a different problem. So, to estimate how much the firings rates change after the problem layout switch s(x_ij_) was calculated on the 20 trials after the problem layout switch (‘test’ between problems), where *μ*_k_ and *σ*_k_ were calculated from the ‘train’ trials from a different layout. This measure was calculated for each neuron separately and then averaged across all neurons for each brain region.

### Singular Value Decomposition

Singular value decomposition (SVD) was performed using the numpy linalg.svd function in Python. SVD is a principal component analysis technique that decomposes any n x m matrix into a product of three matrices:

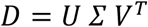

where *D* comprises the data matrix to be decomposed, *U* and *V^T^* are sets of singular vectors capturing patterns of covariation in the data, and *Σ* is a diagonal weight matrix.

In our SVD analyses, each row of *D* was the demeaned, trial-aligned activity of one neuron across each time-point of four concatenated trial types; rewarded A choices, non-rewarded A, rewarded B, and non-rewarded B, so the shape of D was [n_neurons, 4*n_timepoints_per_trial]. The columns of *U* are vectors which we term cellular modes because each is a set of weights over neurons, representing groups of neurons whose activity covaries. Each cellular mode has a corresponding row in *V^T^* which we term a temporal mode, as it is a set of weights over time-points, representing the time-course of the cellular mode’s activity. Each temporal mode spans the same set of time-points as the data matrix, and hence captures variation both over time-in-trial and trial type. As both modes are unit vectors, their contribution to the total data variance is determined by the corresponding element of the diagonal matrix *Σ*.

The cellular modes are given by eigendecomposition of the covariances between neurons, as can be seen from the following:

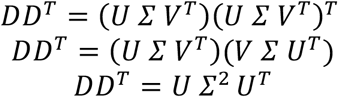

As *DD^T^* is the non-normalised covariance between neurons across time-points, *Σ*^2^ is a diagonal matrix of eigenvalues, *U* are the corresponding eigenvectors, and *U^T^* = *U*^−1^ because *U* is an orthonormal basis.

Similarly, the temporal modes are given by eigendecomposition of the covariances between time-points

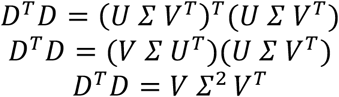

As *D^T^ D ^T^* is the non-normalised covariance between time-points across neurons, *Σ*^2^ is a diagonal matrix of eigenvalues, *V* are the corresponding eigenvectors, and *V^T^* = *V*^−1^ because *V* is an orthonormal basis.

Our goal was to test whether cellular and temporal patterns generalise across different problems, by quantifying how well cellular and/or temporal modes from one problem explained variance in another. As a control for drift in representations over time, we compared generalisation between problems with generalisation to held out data from the same problem. To do this we constructed separate data matrices for the first and second half of each problem:

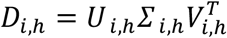

where *i* is the problem number *i* = {1,2,3} and *h* is the half of the problem the data is taken from *h* = {*f,s*}. We can then compare generalisation between the second half of one problem with the first half of the next, with generalisation between first and second half of the same problem, to ensure any drift is matched between within and cross problem comparisons.

We quantified three different ways in which activity patterns might generalise between problems; 1) Generalisation of temporal modes irrespective of whether or not they recruited the same neurons. This corresponds to the same trial events being represented but not necessarily by the same neurons. 2) Generalisation of cellular modes irrespective of whether they have the same time course. This corresponds to the same cell assemblies co-activating but not necessarily representing the same trial events. 3) Generalisation of cellular-temporal mode pairs. This corresponds to the same cell assemblies representing the same trial events across problems.

To quantify how well temporal modes generalised across problems we projected the data matrix from half of one problem on the temporal modes from an adjacent half of a different problem:

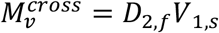

The total variance explained by each temporal mode for this problem pair is given by squaring the elements of 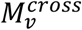 and summing over neurons. We average across all adjacent problem pairs and plot the cumulative variance explained as a function of the number of temporal modes used.

The corresponding within-problem variance explained is given by projecting the data matrix from half of one problem onto the temporal modes from the other half of the same problem:

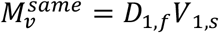

Similarly for the cellular modes, the cross-problem generalisation was given by projecting the data matrix from half of one task on the cellular modes from an adjacent half of a different problem:

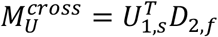

The total variance explained by each cellular mode for this problem pair is given by squaring the elements of 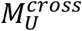 and summing over time-points. Again, we average across all adjacent problem pairs and plot the cumulative variance explained as a function of the number of cellular modes used.

The corresponding within-problem variance explained is given by projecting the data matrix from half of one problem onto the cellular modes from the other half of the same problem:

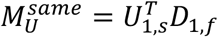

To quantify how well pairs of neural and temporal patterns generalised between problems, we projected the data matrix from half of one problem on the cellular and temporal modes from an adjacent half of a different problem:

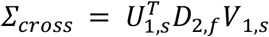

*Σ_cross_* is not diagonal, however if the same cell assemblies perform the same roles in two problems, the temporal and cellular modes will align, and *Σ_cross_* will have high weights on the diagonal. We therefore plotted the cumulative sum of the squared weights of the diagonal elements. Since we had different numbers of neurons in each brain region *Σ_cross_* was normalised by the number of neurons recorded from the respective brain region.

The corresponding within-problem variance explained is given by projecting the data matrix from half of one problem onto the cellular and temporal modes from the other half of the same problem:

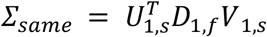

To determine the significance of the differences between two regions we compared differences in the data between PFC and CA1 against a null distribution of differences between areas under the curve by shuffling the sessions between CA1 and PFC animals.

### Estimating Policy

To estimate policy, we first fit a logistic regression predicting current choices with a history of choices, rewards, and choice x reward interactions across all sessions (Figure 5A). To calculate the effect of these variables on an animal’s choice (policy) on individual trials we then computed a product of these average coefficients with the animal’s history of rewards, and choice x reward on individual trials on each session. This gave one number on each trial describing the probability that the animal would choose A given its previous choices and rewards (policy). We then performed a cross-trial regression using this policy estimate and its interaction with current choice (chosen policy) together with other behavioural variables (choice, outcome, and outcome x choice interaction on the current trial) to establish unique variance explained (CPDs) by animal’s policy.

To understand whether this signal generalises between different problems and locations we next conducted this analysis independently on A and B choices (here only able to predict firing rates from only reward and policy). We compared the coefficients from these models across problems by computing correlations across all time points (Figure 5C). Finally, to understand whether these across trial policy signals might also be tied to representations of unique trial stages we examined time-slices through the correlation matrices at initiation, choice, and outcome times. The differences between these signals at each timepoint were then compared against null distributions described in the Statistical Significance Methods section.

### Decoding Analyses

To show a categorical difference between the features encoded by the two regions we adopted a decoding analysis based on support vector classification (sklearn.svm.SVC). Specifically, we trained the decoder to classify different stages of the trial (Initiation, Choice A, *Choice* B, Choice A Reward, Choice A No Reward, Choice B Reward, Choice B No Reward) on *one* of the problems and tested how well it performed on classifying these same trial stages in a *different* problem. This was computed for all problem pairs and the mean decoding accuracy for each trial stage was shown in a form of a confusion matrix.

We next summed the decoding probabilities that are associated with a representation of (i) physical port, (ii) trial stage (initiation, choice, type of outcome) and (iii) abstract choice. Because in 1 of our problem layout pairs initiation port became a B choice (Layout 2 to Layout 3) and in another initiation became a B choice (Layout 3 to Layout 2) mistakes made by the decoder between B choice and Initiation in these layout pairs highlight a prominent representation of port location. The mistakes made between A choices and B choices, A rewards and B rewards and A no-rewards and B no-rewards highlight a representation of trial stage. Lastly, representation of an abstract B choice was computed by summing the lower half of the diagonal that represents the decoding accuracy related to decoding the same abstract choice but in a different physical location across problems. The statistical significance of the differences in decoding of these features was established by permuting animal identities between regions and comparing the real differences between regions against the 95% confidence interval of the shuffle.

### Additional Controls for Physical Movement

To provide additional controls for movement related activity we sought to eliminate the effect of animal’s position, velocity, and acceleration on firing rates before performing all our subsequent analyses. To do this we used DeepLabCut^66^ pose estimation software to extract the animal’s nose position in each session. Because the cameras in the operant boxes were located above the animal there were some artefacts in the tracking caused by the occlusion of the nose by the ports and the head-cap, which caused the estimated position to jump to incorrect locations. To correct for this we first removed all samples where the likelihood of correct estimation output by DLC was below 90%. We then removed samples adjacent to jumps in position larger than 10 times the standard deviation of displacements between frames, estimated using the 16th and 84th percentiles of the displacement distribution. We then removed samples that were not in contiguous groups of at least 5. After this artefact removal step we interpolated the missing data, taking advantage of the fact that the movements of the ears and nose are highly correlated, such that the trajectories of the ears provide information about movements of the nose when the nose is occluded. The interpolation was implemented by minimising a cost function with two terms: i) the sum of squared derivatives of the nose position – which promotes linear interpolation of missing data and ii) the sum of squared differences between the derivatives of the ear and nose positions – which promotes the interpolated trajectory of the nose tracking those of the ears.

Next, since we had the ground truth of our port locations in physical space, we performed a linear registration and transformed the 2D coordinates extracted from the video from the oblique camera view to a more informative horizontal view of the wall of the ports. Finally, we utilised our behavioural data to find when the animals were inside the ports, and corrected for any inaccuracy in our DLC data by placing these coordinates inside the ports.

As we did not expect that the 2D coordinates of animal’s nose position would be linearly related to neural firing rates (e.g., due to previously reported existence of ‘place cells’ in CA1) we first needed to create vectorised ‘occupancy maps’ (Supplementary Fig 7A). Specifically, we defined a set of Gaussian “radial basis functions” with the centres randomly selected from an animal’s 2D coordinates in each session and a standard deviation of 1 cm. Next, for each time point we calculated the activity of each basis function (Gaussian in distance from centre of this field), resulting in a matrix of shape [time, n_basis_functions].

To account for cross-correlations in this matrix we next did a principal component analysis to extract the first ten orthogonal occupancy components across time accounting for 95> % of variance. To confirm that our key results could not be explained by movement related parameters, we repeated our main analyses using the residual firing rates from a linear model predicting the firing of each neuron using these occupancy components, as well as the acceleration and velocity of the animal at each timepoint (Supplementary Figure 7). Because we did not have video data for all our animals due to technical limitations at the time of experiments the significance between brain areas in these analyses was only computed by shuffling the sessions of CA1 and PFC animals to obtain null distributions and correcting for multiple comparisons as before (see *Statistical Significance Methods*).

