## Supplementary Figures for "Complementary task representations in hippocampus and prefrontal cortex for generalising the structure of problems"

### 4 – Supplementary Figures

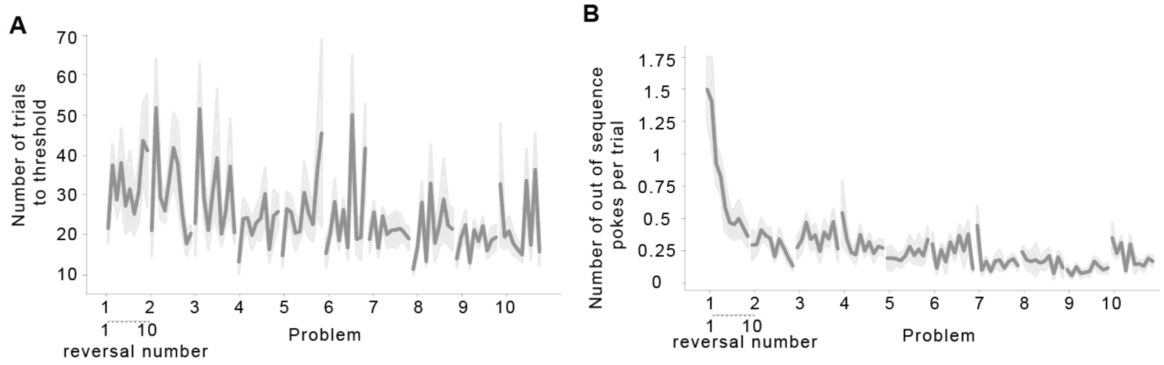


**Supplementary Figure: 1. Transfer learning in mice. A)** Number of trials following a reversal taken to reach the threshold to trigger the next reversal, as a function of reversal number within each problem and problem number. **B)** Number of pokes per trial to a choice port that was no longer available because the subject had already chosen the other port, as a function of reversal number within each problem and problem number. Error bars report the mean ± SEM across different mice.

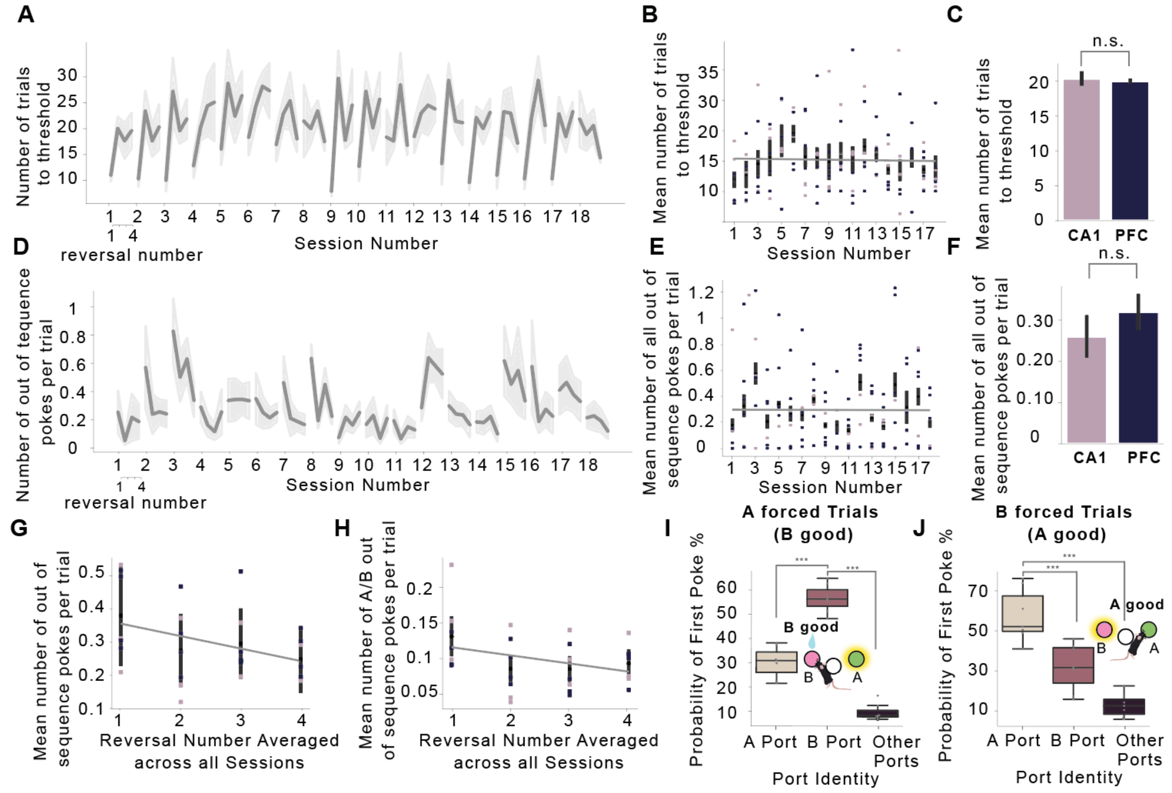


**Supplementary Figure: 2. Behaviour during recordings. A)** Number of trials following a reversal taken to reach the threshold to trigger the next reversal, as a function of reversal number within each session ($F_{(3,18)} = 27.6$, $p < .001$) and session number ($F_{(17,102)} = 0.86$, $p = .62$) during recordings. **B)** Average number of trials following a reversal taken to reach the threshold to trigger the next reversal, as a function of session number during recordings (analogous to Figure 1 E). **C)** There was no significant difference in the mean number of trials animals took to reach the threshold for a reversal during recordings between PFC and CA1 animals ($t_{(7)} = 0.31$, $p = .766$). **D)** Number of pokes per trial to a choice port that was no longer available because the subject had already chosen the other port, as a function of reversal number within each session ($F_{(3,18)} = 8.52$, $p < .001$) and session number ($F_{(17,102)} = 0.86$, $p = .621$) during recordings. **E)** Average number of out of sequence pokes mice made as a function of problem number during recordings (analogous to Figure 1G). **F)** There was no significant difference in the mean number of out of sequence pokes during recordings between PFC and CA1 animals ($t_{(7)} = 0.75$, $p = .489$). **G - H)** Mice made more out of sequence pokes per trial during the first reversal block compared to every following reversal (Reversal 1 vs Reversal 2: $t_{(125)} = 2.54$, $p = .024$), Reversal 1 vs Reversal 3: $t_{(125)} = 3.52$, $p = .004$; Reversal 1 vs Reversal 4: $t_{(125)} = 3.30$, $p = .004$). **G)** All out of sequence pokes during that problem were plotted as a function of each reversal. **H)** For comparison with analogous plot during the behavioural training only the A/B out of sequence pokes during that problem were plotted as a function of each reversal. **I- J)** Mice did not follow lights to complete a trial. **I)** On forced A trials where A choice was illuminated but B choice was good animals were more likely to first choose the B choice port than the A port ($t_{(7)} = 6.44$, $p < .001$) or other choice ports ($t_{(7)} = 8.07$, $p < .001$). **J)** On forced B trials where B choice was illuminated but A choice was good animals were more likely to first choose the A choice port than the B port ($t_{(7)} = 2.70$, $p = .035$) or other choice ports ($t_{(7)} = 3.22$, $p = .018$). Error bars report the mean ± SEM (A-I) or median, inter quartile range and min and max(I-J) across different mice.

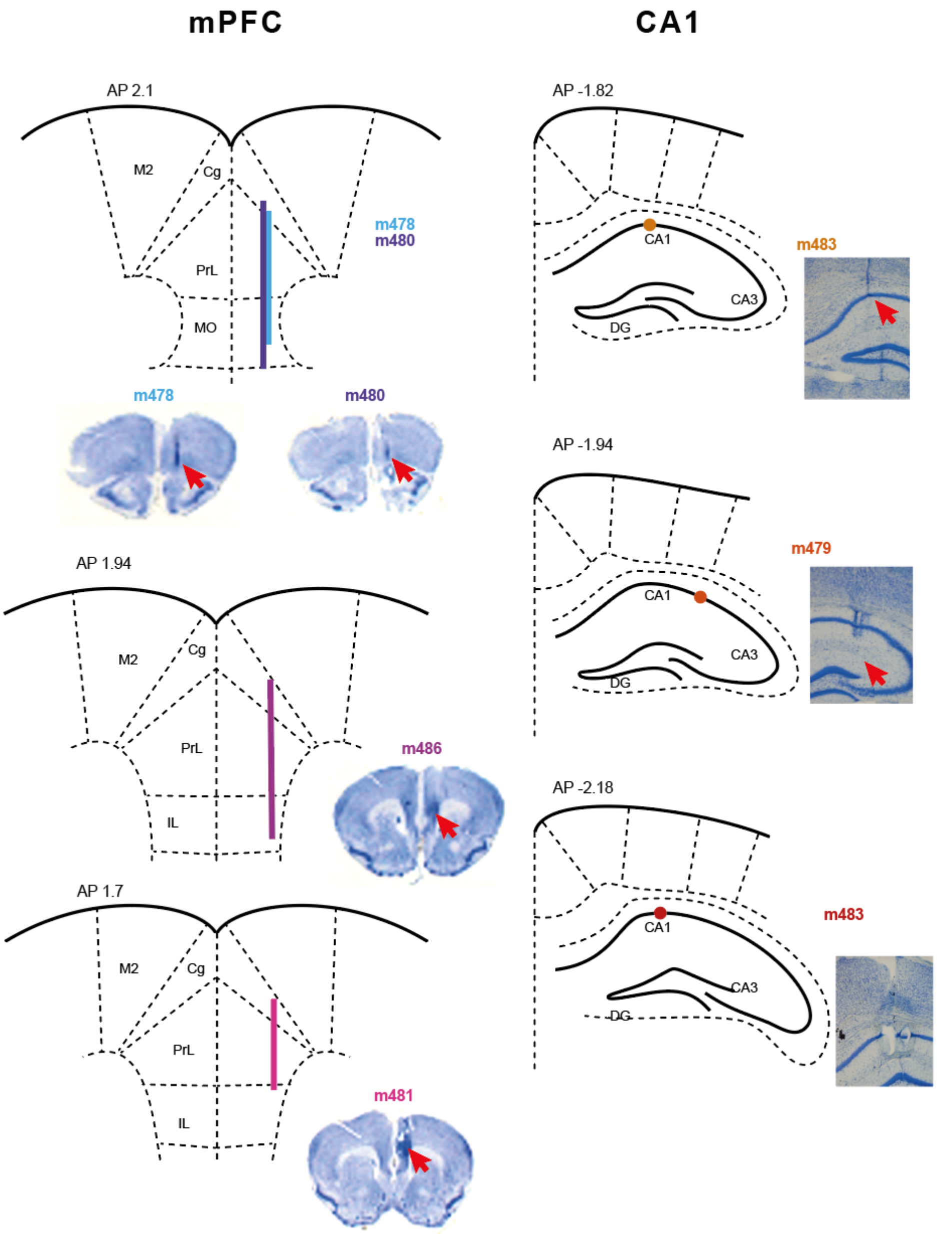


**Supplementary Figure: 3. Location of Recording Sites.** mPFC probes were advanced on each session whereas CA1 probes were static throughout recordings (for more details see *Electrophysiological Recordings and Spike Sorting Methods*). mPFC recordings were mostly from prelimbic area but also in some cases reached infralimbic area, medial orbital and cingulate**.**

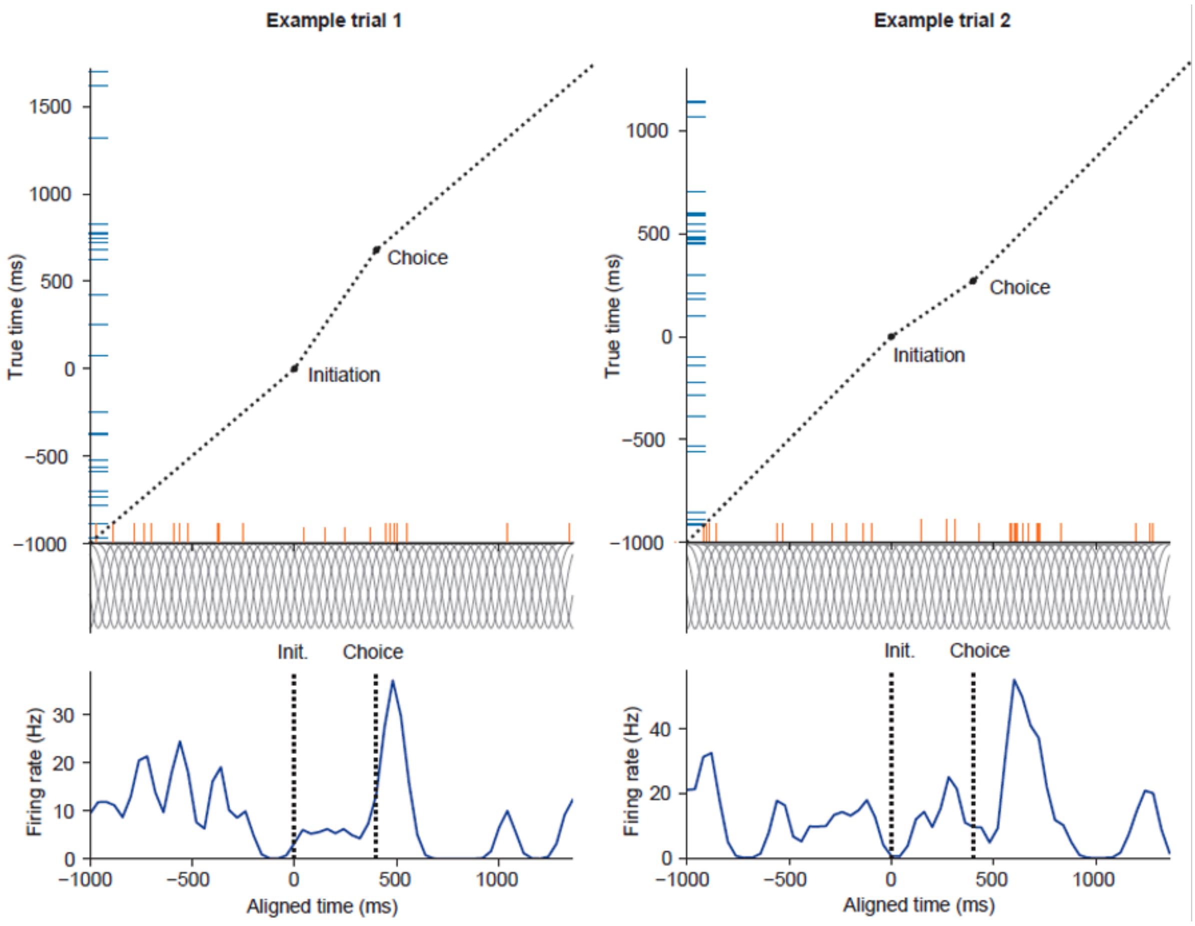


**Supplementary Figure: 4. Trial alignment of activity.** Diagram illustrating alignment of spike activity across trials. Trials were aligned using the times of initiation and choice port entry, by warping the interval between these two events to match the median interval. Top panels show spike times in the true (blue ticks) and aligned (orange ticks) time reference frames. Spike times were transformed by linear interpolation between the reference points. The output firing rate (bottom panels) was calculated at points spaced every 40ms in the aligned reference frame using Gaussian smoothing (40ms standard deviation) of the spike train. To compensate for the change in spike density due to the time warping, spikes were weighed by the stretch factor between the true and aligned reference frames (weighting is indicated by height of the orange ticks) prior to Gaussian smoothing.

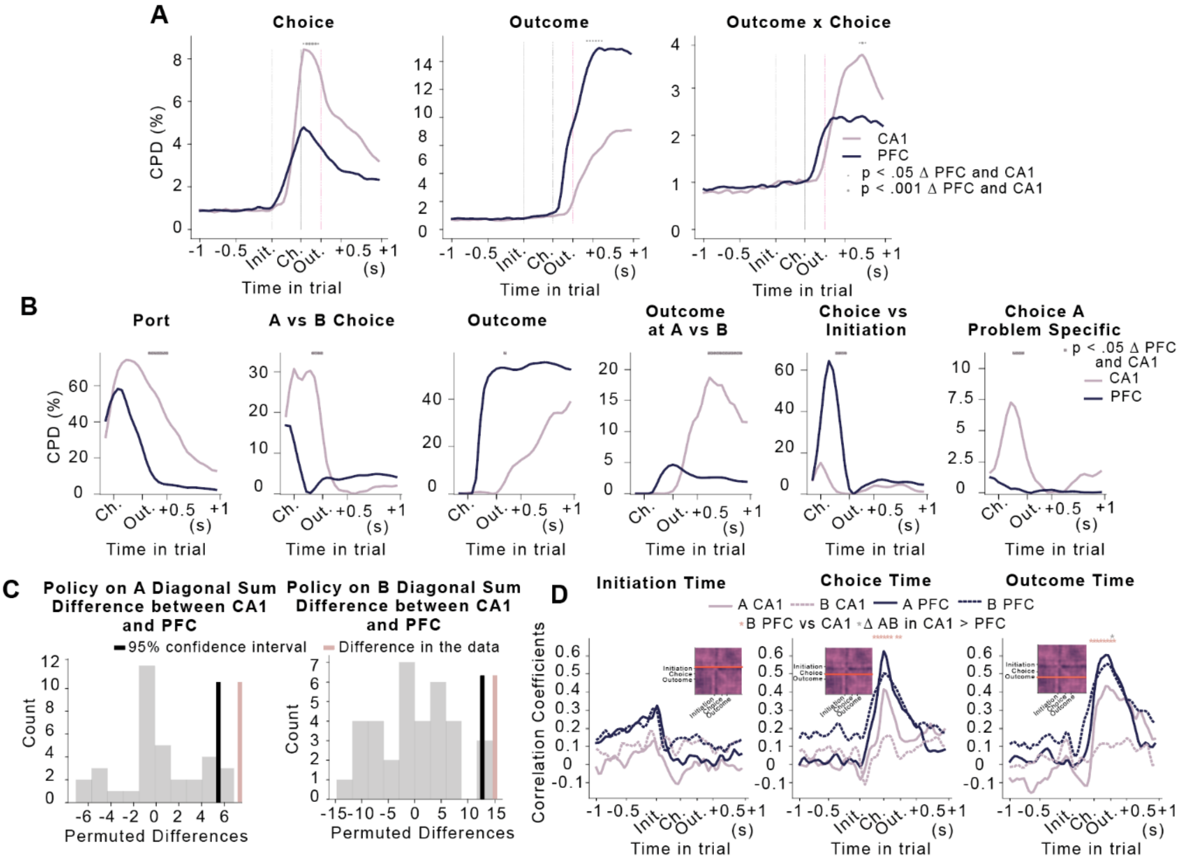


**Supplementary Figure: 5. Replicating main findings of trial stage (A-B) and policy (C-D) representations generalising across different sensory problems more in PFC than CA1 with significance levels based on a group test obtained by shuffling animal identities between regions. A)** Coefficients of partial determination from the linear model shown in Figure 3A for choice, outcome, and outcome x choice regressors in PFC and CA1. **E)** Coefficients of partial determination in a regression analysis modelling the pattern of representation similarities using the RDMs shown in Figure 3D. **C)** Sums along the diagonal of the correlation matrices shown in Figure 5C separately for A and B choices. **E)** Slices through the correlation matrices at initiation (left), choice (centre) and outcome (right) times for A (solid) and B (dash line) choices. For animal shuffles in singular value decomposition analyses see Supplementary Figure 7.

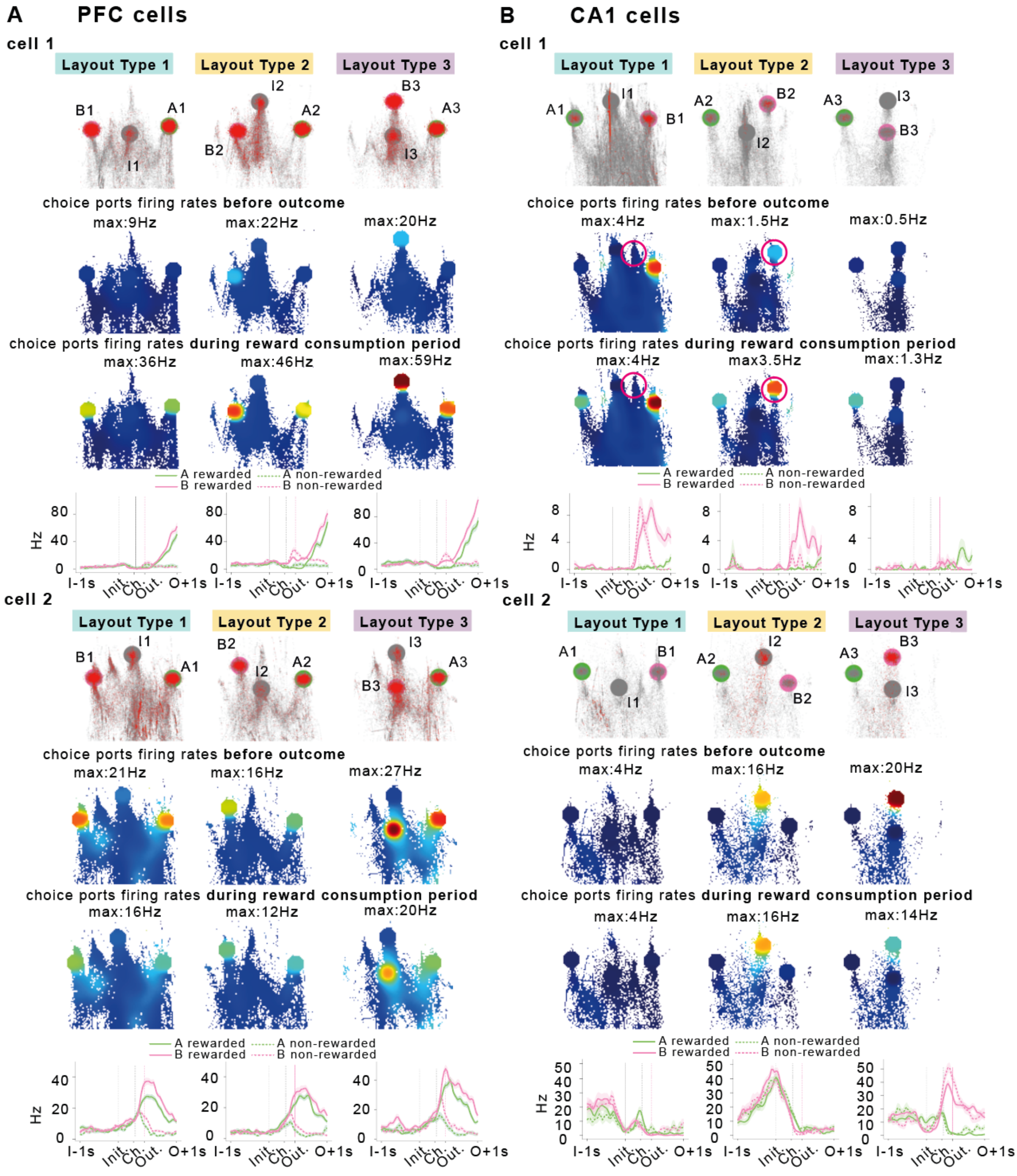


**Supplementary Figure 6. Additional example PFC and CA1 neurons in physical space and behavioural task.** For every cell, the top subpanels show nose trajectories in grey and spikes in red in each problem layout, in a 2D space corresponding to the view of a camera positioned above the box looking at the ports, affine transformed to correct for the oblique view of the ports, (initiation port is indicated in grey, A ports in green and B ports in pink). Middle panels show firing rate heat maps,plotted either including spikes and occupancies in choice ports before reward is delivered, or during reward consumption. Bottom panels show corresponding task event aligned activity. **A) PFC cells**. Cell 1 is a reward cell and fires at all choice ports during the reward consumption period. Cell 2 is a rewarded choice cell and starts to fire for all rewarded choices before the reward is released. **B) CA1 cells.** Cell 1 has a conjunction of space and reward, only firing for B-rewards at ports in the upper right portion of the map. Note that in layout 1 the animal does in fact make some error pokes into a port that is inactive, but will be the B port in layout 2 (highlighted with a circle). It fires at this port in layout 2 (where it gets a reward) but not layout 1. Cell 2 is a port selective cell that always fires at the same port, no matter whether it is choice or initiation. These types of cells will be accounted for by our controls for physical movements and precise 2D nose position (see Supplementary Figure 7 and *Additional Controls for Physical Movement Methods*).

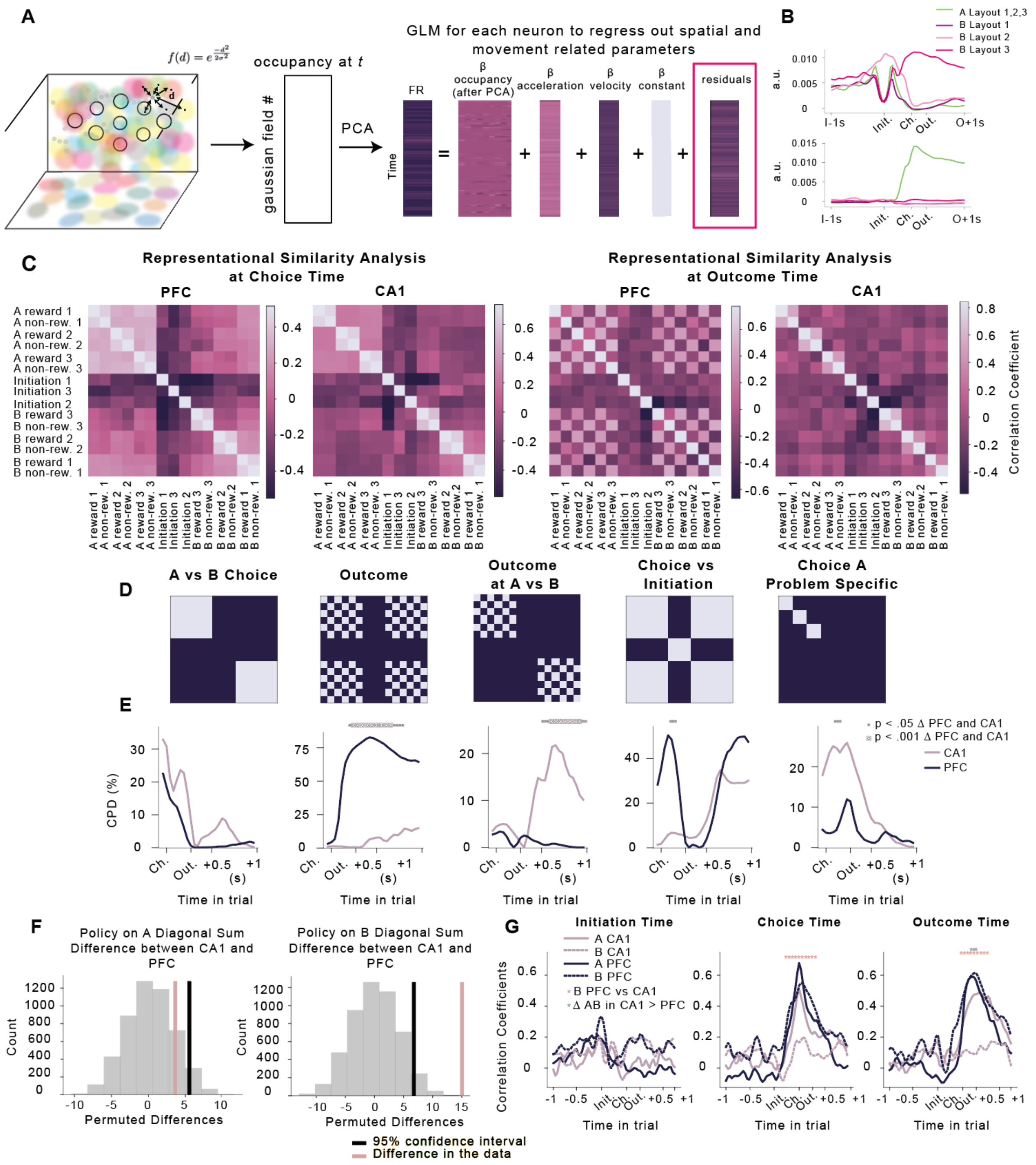


**Supplementary Figure: 7. Replication of main findings after accounting for fine-grained movement related activity**. **A)** We do not expect 2D coordinates to be linearly related to firing rates so to account for place cell like coding of nose position in the firing rates of neurons we defined a set of gaussian "radial basis functions" with the centres randomly selected from an animal's 2D coordinates in each session (different coloured circles). Next, for each time point we calculated the activity of each basis function (gaussian in distance from centre of this field) resulting in a time x (# of basis functions) matrix (left). To account for cross-correlations in this matrix we next did a principal component analysis to extract the first ten orthogonal occupancy components across time (middle). Next, we fit a linear regression model predicting firing rates of neurons with occupancy as well as velocity, and acceleration predictors resulting in residual firing rates that do not contain variability related to these movement related parameters (right). **B) Top two principal components from the PCA analysis of occupancies in A from an example session.** The first component differentiates initiation in Layout 1 and port B in Layout 2 (same physical location) from other ports. The second component differentiates port A (same physical location) from all other ports. **C)** Representation similarity at 'choice time' (left) and 'outcome time' (right), quantified as the Pearson correlation between the **residual** neural activity (after accounting for movement related parameters) vectors for each pair of task conditions as in Figure 4C. **D)** Representational Similarity Design Matrices (RDMs) used to model the patterns of representation similarity observed in the data. Port RDM was not included in this analysis as PCs we regress out to account for movement related activity correlate strongly with the port position in the task (as in **B**). Using an RDM that is so highly correlated with a parameter that has already been regressed out can lead to false correlations (a la Berkson's paradox). **E)** Coefficients of partial determination in a regression analysis modelling the pattern of representation similarities in residual firing rates using the RDMs in D. **F-G)** Policy analyses conducted on residual firing rates after accounting for movement related activity. The generalisation of policy on A trials is no longer stronger in PFC than CA1. Controls of precise physical movements in singular value decomposition analyses are presented in the main text (Figure 4D-J).

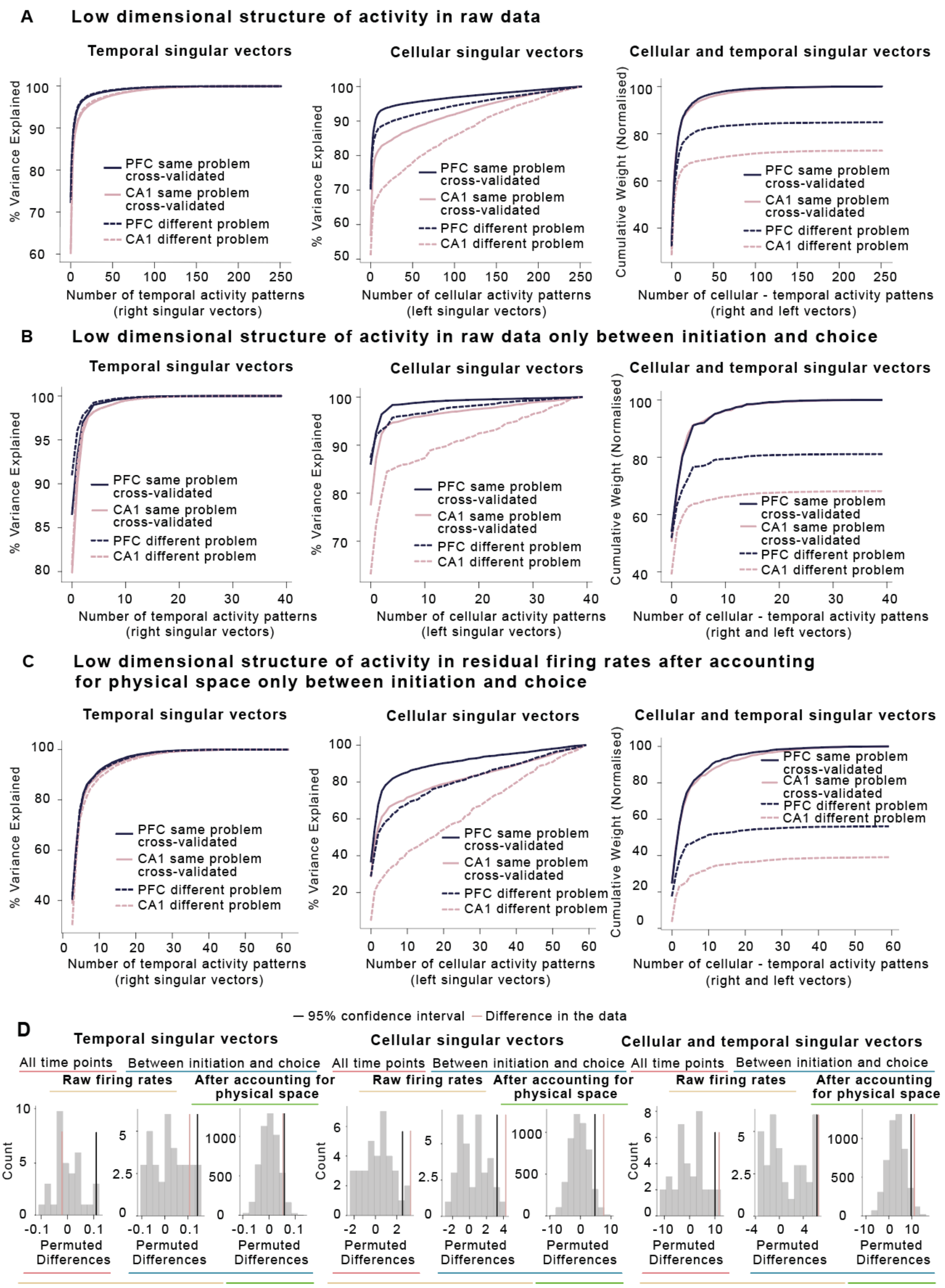


**Supplementary Figure: 8. Additional analyses of low dimensional structure of activity in PFC and CA1. A – C)** Left: Variance explained when using temporal activity patterns $V_I^T$ from one problem to predict either held out activity from the same problem (solid lines) or activity from a different problem (dash lines). Middle: Variance explained when using cellular activity patterns $U_I$ from one problem to predict either held out activity from the same problem (solid lines) or activity from a different problem (dash lines). Right: Cumulative weights along the diagonal $\sum$ using pairs of temporal $V_I^T$ and cellular $U_I$ activity patterns from one problem to predict either held out activity from the same problem (solid lines) or activity from a different problem (dash lines). **A, B)** Low dimensional structure of activity in raw data across all time points in the trial **(A)** and only taking activity between initiation and choice **(B)**. **C)** Low dimensional structure of activity after accounting for movement related parameters in the data and only running the analyses on time points between initiation and choice. **D)** Permutation tests of temporal, cellular and cellular and temporal singular vector differences based on the null distribution obtained by shuffling animals across groups (raw firing rates) or sessions (residual firing rates after accounting for physical space). We could not run permute animals in the analyses of residual firing rates because we were not set up for recording video data for our first implanted animal.

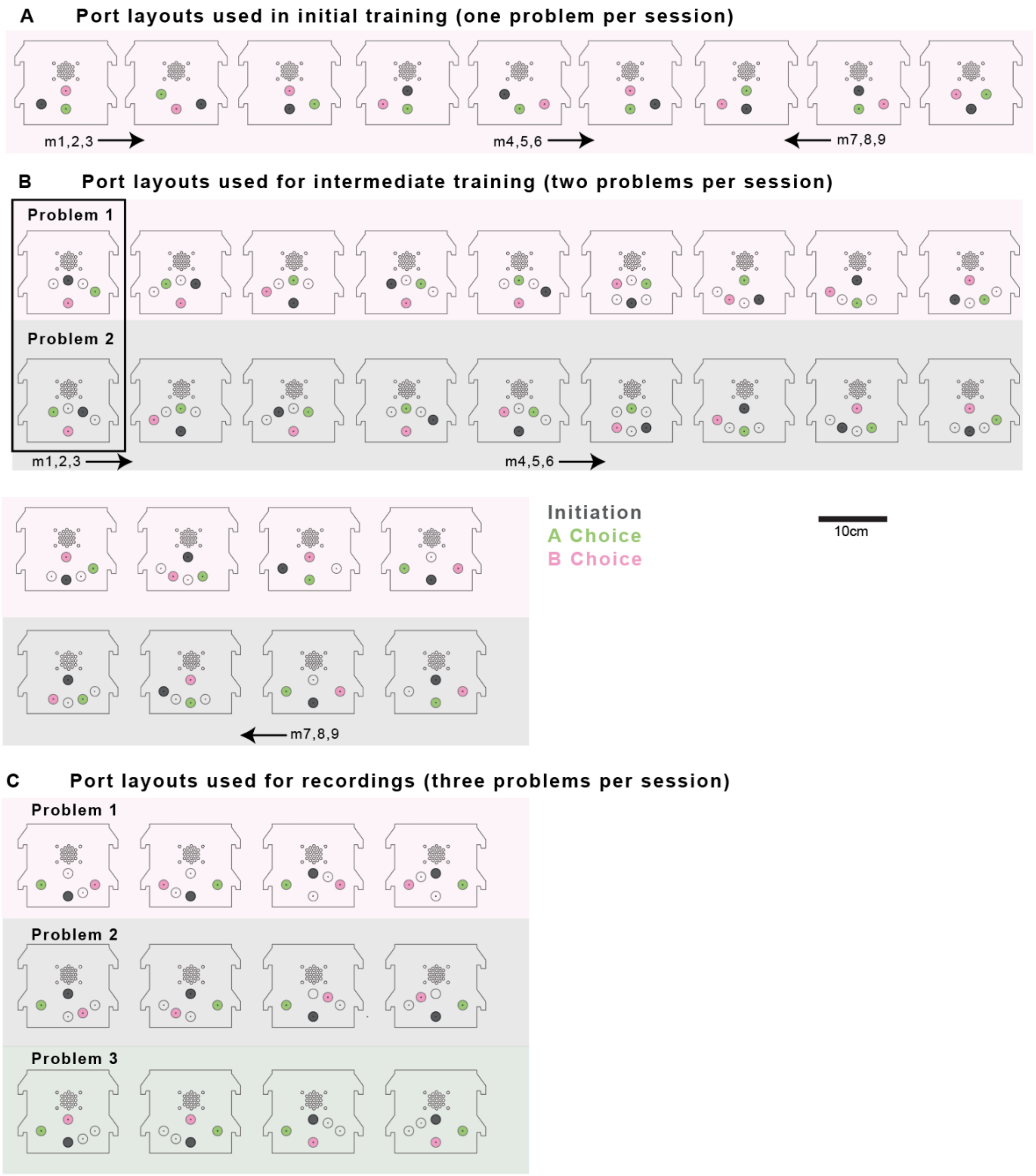


**Supplementary Figure: 9. Counterbalancing and port layouts used throughout the experiments.** Port layouts used for initial training (A), intermediate training (B) and recording sessions (C), showing the locations of the initiation (grey), A choice (green) and B choice (pink ports) on the wall of the box. Ports that were not used in any of the problems presented in a session were covered up and are not shown on the diagrams. Ports that were exposed but not used in a given problem are shown in white. In B and C the set of problems used in a single session are arranged vertically. During initial and intermediate training stages, the presentation order of different configurations was counterbalanced across animals by randomly assigning mice to 3 in groups each of 3 animals. The starting configurations for the different animals are indicated under the layouts, with an arrow indicating the direction the group subsequently progressed through the layouts across training. The layouts shown for the recordings are labelled 1,2 and 3 for consistency with how they are described in the results text, but the actual presentation order within each session was randomised. For more details on the training protocol see *Behavioural Training Methods.*

.

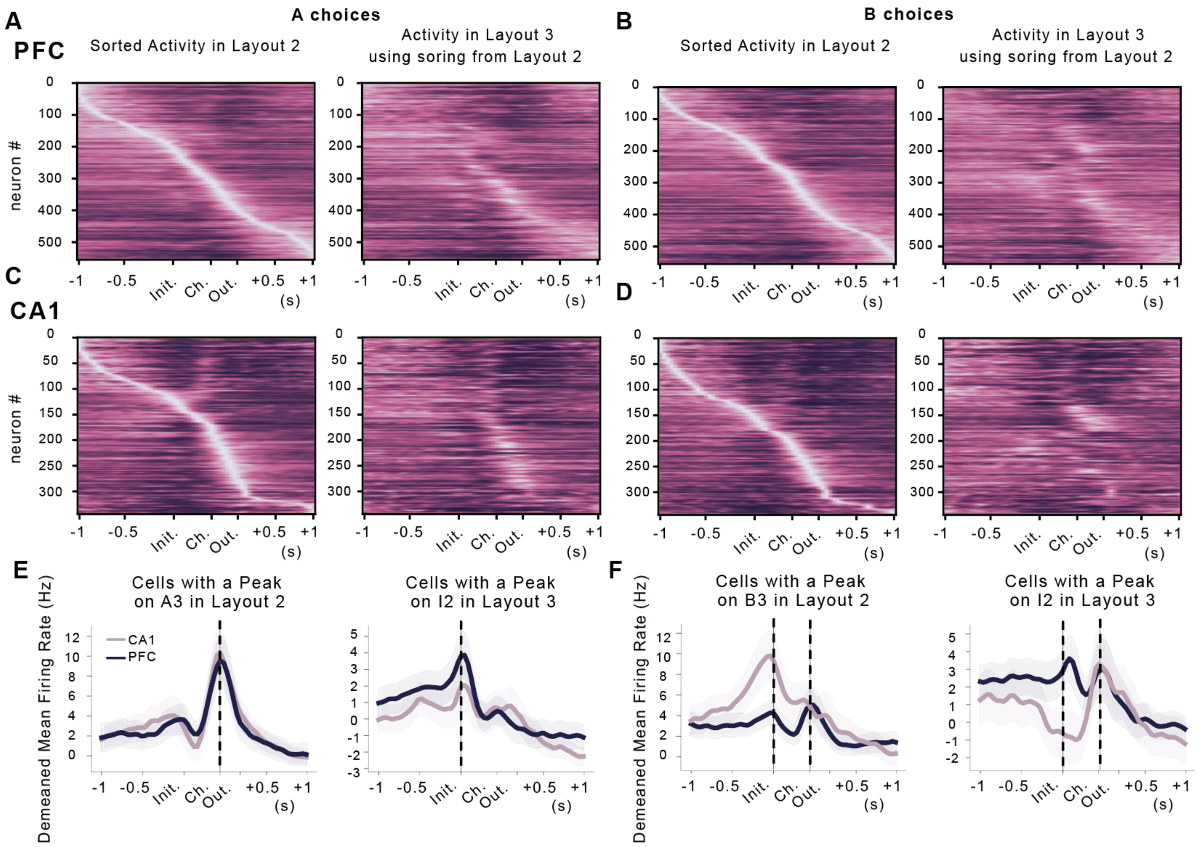


**Supplementary Figure: 10. Port selectivity is more pronounced in CA1 than PFC. A-D)** To evaluate the relative effects of conflict between problem general and port specific representations we sorted neural activity in Layout Type 2 and used this sorting to plot the activity in Layout Type 3. Choice B in Problem Layout 2 is Initiation in Problem Layout 3 but for comparison we also plotted A choices that were in the same physical port. **A)** PFC sorted activity in trials where A port was chosen in Layout Type 2 (left) and activity in trials where A port was chosen in Layout Type 3 using the sorting from Layout 2. **B)** PFC sorted activity in trials where B port was chosen in Layout Type 2 (left) and activity in trials where A port was chosen in Layout Type 3 using the sorting from Layout 2. **C-D)** Same as **A-B** but for CA1. **E, F)** We identified cells that had their peak firing rates around initiation and choice events in one layout type (A-D left) and plotted the average activity of these cells in the other layout. **E-F)** Average activity of cells in Layout 2 that had a peak at A choice time in Layout 3. **E)** Average activity of cells on A choices in Layout 2 that had a peak at A choice time in Layout 3 (left). Average activity of cells on A choices in Layout 3 that had a peak at initiation time in Layout 3 (right). **F)** Average activity of cells on B choices in Layout 2 that had a peak at B choice time in Layout 3 (left). Average activity of cells in B choices Layout 3 that had a peak at initiation time in Layout 2 (right). CA1 neurons were selective to physical location rather than trial stage. CA1 neurons that responded to initiation (I2) in one problem responded to B choice (B3) in another problem when the I2 became B3 (**F** right) and vice versa (choice B3 neurons in problem layout 3 were selective to I2 in problem 2, **F** left). PFC neurons that peaked at initiation time (I2) in one problem generalised to initiation time (I3) in another problem but also had a peak at choice (B3) time (**F** right) and vice versa (choice B3 neurons also had two peaks at I2 and B2, **F** left).

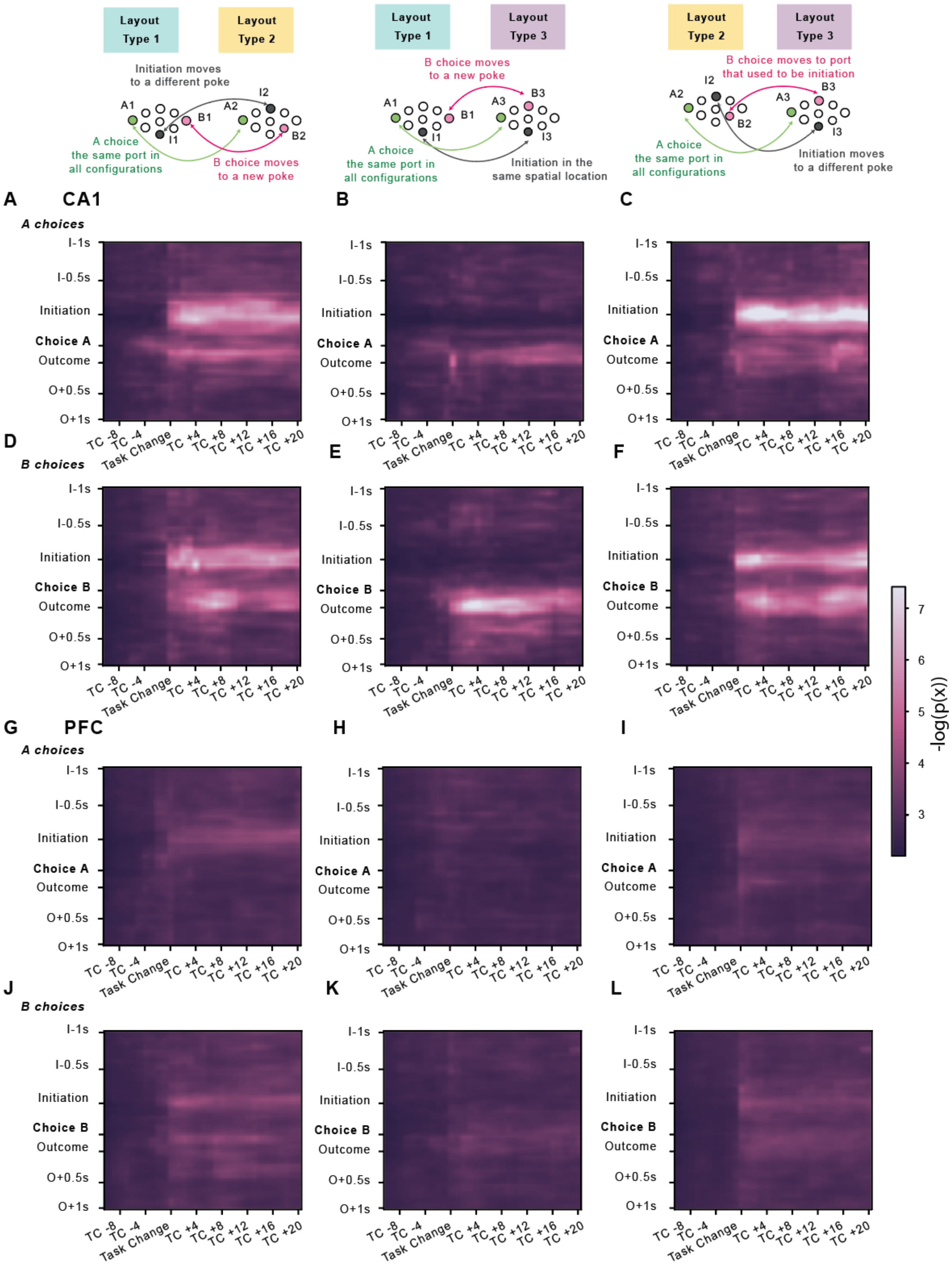


**Supplementary Figure: 11. Rapid problem-induced 'remapping' in CA1 but not PFC.** Diagram showing how initiation and choice ports changed position for three types of problem layout transition analysed in the left, middle and right columns of the figure. 'Remapping' quantified by 'surprise' (for more details of the analysis see *Surprise Measure Methods*) was computed between three types of problem switches. In the first type of switch (left) both initiation and B choice were in different locations in two problems. In the second type (middle) initiation was in the same physical location but B choices were in different ports. In the third type (right) initiation was in different physical locations but initiation port in layout 2 was in the same location as choice B in layout 3. A choice port was always in the same physical location in all problem layouts. **A-J)** Heatmaps showing how surprising the activity at each time point of each trial around a layout transition was, with respect to the distribution of activity at the same time point in a ‘baseline’ period just prior to the plotted trials. Activity in CA1 on A and B choice trials is shown in **A-C** and **D-F** respectively. Activity in PFC on A and B choice trials is shown in **G-I** and **J-L** respectively. In CA1, when an initiation or choice port moved to a different physical location, the neuronal representation at the corresponding stage of the trial changed immediately, as indicated by an abrupt increase in surprise at the layout transition **A-F**.
